# Development of white matter tracts between and within the dorsal and ventral streams

**DOI:** 10.1101/2021.01.27.428423

**Authors:** S. Vinci-Booher, B. Caron, D. Bullock, K. James, F. Pestilli

**Author notes:** **Corresponding Authors**: Franco Pestilli, Department of Psychology, The University of Texas at Austin, 108 E Dean Keeton St, Austin, TX 78712, (347) 829- 4185, Sophia Vinci-Booher, Department of Psychological and Brain Sciences, Indiana University, 1101 E. 10th Street, Bloomington, IN 47405. **Author Contribution Statement** Sophia Vinci-Booher contributed to all aspects of the manuscript, including the original conception of the study, ongoing conceptual development, the design, data collection, analyses and software, writing the original draft of the paper, and revisions. Brad Caron and Dan Bullock contributed software and supported software development, participated in data quality checks, and commented on the manuscript. Karin James contributed to data collection, and commented on the manuscript. Franco Pestilli contributed to the original conception of the study, the conceptual development of the work, the design, the analyses, software, training of Sophia Vinci-Booher, and in the writing of the manuscript and revisions.

## Abstract

The degree of interaction between the ventral and dorsal visual streams has been discussed in multiple scientific domains for decades. Recently, several white matter tracts that directly connect cortical regions associated with the dorsal and ventral streams have become possible to study due to advancements in automated and reproducible methods. The developmental trajectory of this set of tracts, here referred to as the posterior vertical pathway (PVP), has yet to be described. We propose an input-driven model of white matter development and provide evidence for the model by focusing on the development of the PVP. We used reproducible, cloud-computing methods and diffusion imaging from adults and children (ages 5-8 years) to compare PVP development to that of tracts within the ventral and dorsal pathways. PVP microstructure was more adult-like than dorsal stream microstructure, but less adult-like than ventral stream microstructure. Additionally, PVP microstructure was more similar to the microstructure of the ventral than the dorsal stream and was predicted by performance on a perceptual task in children. Overall, results suggest a potential role for the PVP in the development of the dorsal visual stream that may be related to its ability to facilitate interactions between ventral and dorsal streams during learning. Our results are consistent with the proposed model, suggesting that the microstructural development of major white matter pathways is related, at least in part, to the propagation of sensory information within the visual system.

**Significance Statement:** Understanding white matter development is important to building predictive models that can inform interventions and targeted educational methods. We propose and provide evidence for an input-driven model of white matter development. We tested an uncharacterized aspect of human brain development. Namely, how the recently described posterior vertical white matter tracts develop. Our results suggest a developmental progression along the known, direct anatomical connections from posterior visual areas to anterior ventral and dorsal areas. Our results suggest fundamental biological mechanisms that clarify the role of white matter in predicting human learning and behavior.

## Introduction

Visual information in the human brain is processed along two processing streams, the dorsal and ventral streams (M. A. Goodale & Milner, 1992; Milner, 2017; Milner & Goodale, 2008; Saber et al., 2015; Sani et al., 2019; Takemura et al., 2015; Ungerleider & Haxby, 1994). A large body of work has demonstrated a degree of functional segregation between perceptual processing associated with the ventral stream and processing for action associated with the dorsal stream (Culham et al., 2003; M. A. Goodale & Milner, 1992; M. Goodale & Milner, 2013; K. H. James et al., 2001; T. W. James et al., 2003; Milner & Goodale, 2008; Ungerleider & Haxby, 1994). However, these two streams do interact (Janssen et al., 2018; Mahon et al., 2013; Milner, 2017; Saber et al., 2015) and a handful of studies suggest that the two streams interact differently in childhood than in adulthood (Freud et al., 2019; Hanisch et al., 2001).

Recent work has reported a set of vertical white matter tracts that directly connect cortical regions associated with ventral and dorsal streams (D. Bullock et al., 2019; Catani et al., 2005; A. Kamali et al., 2014; N. Makris et al., 2013; Nikos Makris et al., 2009, 2017; Maldonado et al., 2013; Menjot de Champfleur et al., 2013; Takemura et al., 2015; Weiner et al., 2017; Wu et al., 2016). We refer to this collection of white matter tracts as the Posterior Vertical Pathway (PVP; see **Figure 1a**). The PVP comprises four tracts for which automated segmentation algorithms have only recently been made available (D. Bullock et al., 2019). Two of these tracts connect posterior ventral temporal cortex to either inferior parietal cortex (i.e., posterior Arcuate, pArc) or superior parietal cortex (i.e., Temporal Parietal connection to the Superior Parietal Lobe, TP-SPL), while the other two tracts connect anterior ventral temporal cortex to either inferior parietal cortex (i.e., Middle Longitudinal Fasciculus to the Angular Gyrus, MDLFang) or superior parietal cortex (i.e., Middle Longitudinal Fasciculus to the Superior Parietal Lobe, MDLFspl). Understanding the development of the PVP can help clarify the mechanisms underlying interactions between the dorsal and ventral streams in childhood and in adulthood.

**Figure 1.**
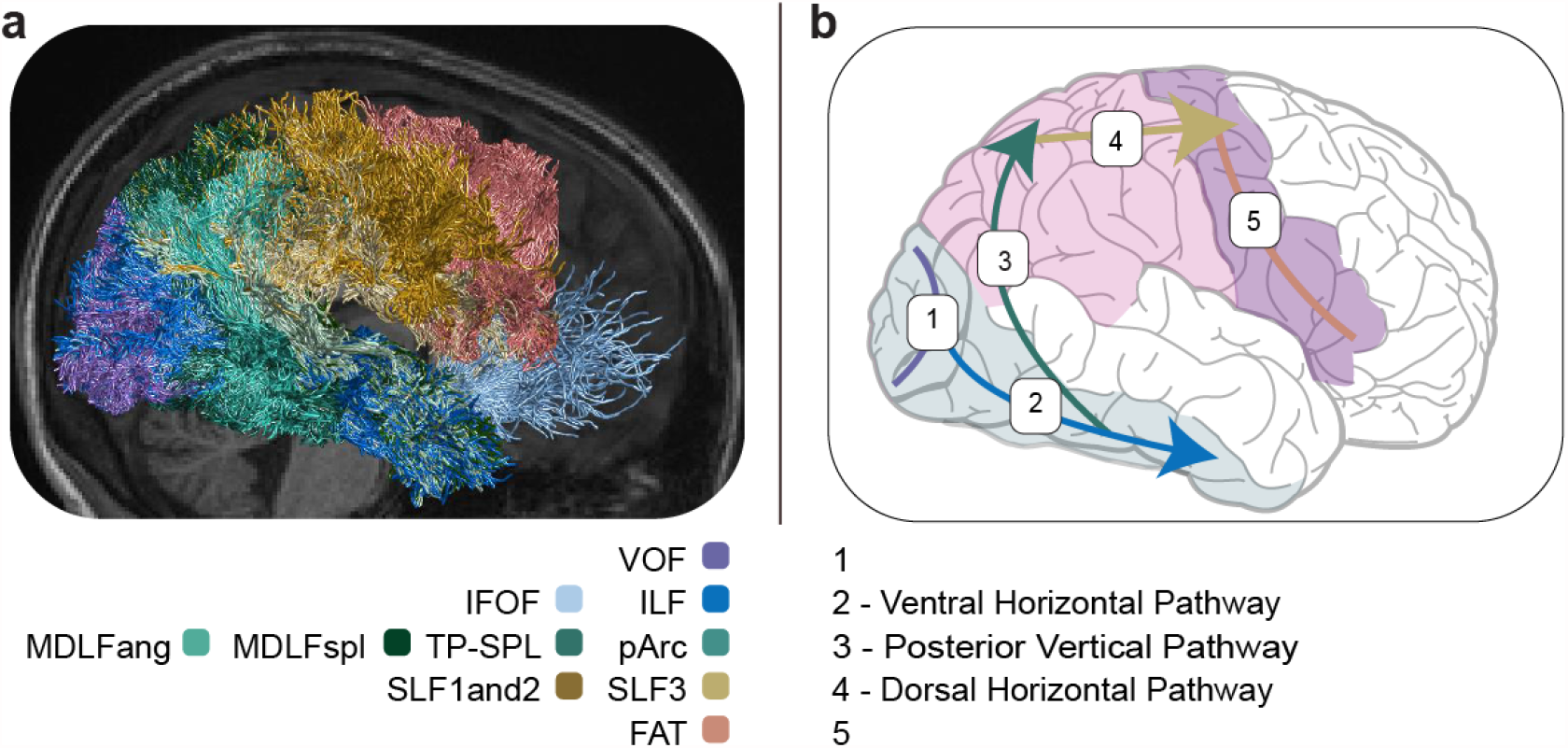
The input-driven model of white matter development. The input-driven model of white matter development posits that some portion of the development of white matter tracts is based on receiving input. **a. White matter tracts of interest displayed on an adult brain**. Tracts included in the ventral horizontal pathway (VHP; blues) included the ILF and IFOF. Tracts included in the posterior vertical pathway (PVP; greens) included the pArc, TP-SPL, MDLFspl, and MDLFang. Tracts included in the dorsal horizontal pathway (DHP; yellows) include SLF1and2 and SLF3. The VOF (purple) and FAT (salmon) were tracts, not pathways, that we analyzed individually as representatives of the proposed earliest and latest developing tracts. **b. Developmental progression predicted by an input-driven model of white matter development**.The model predicts that white matter tracts earlier in the visual hierarchy would develop earlier than white matter later in the hierarchy (Felleman & Van Essen, 1991; Hegdé & Felleman, 2007; Lerner et al., 2001; Poggio & Ullman, 2013; Riesenhuber & Poggio, 1999; Serre et al., 2007). Anatomical tract tracing studies (Jeremy D. Schmahmann, 2006), developmental work on the horizontal white matter tracts (C. Lebel et al., 2008; Loenneker et al., 2011; Peters et al., 2014; Reynolds et al., 2019; Uda et al., 2015; Yeatman, Dougherty, Myall, et al., 2012), and recent connectivity work (Choi et al., 2020), together, suggest that: Visual information first propagates through the occipital cortex (VOF) and (assuming no direct major white matter tract connecting the occipital to the parietal cortex; see **Introduction**), travels to the ventral temporal cortex. PVP tracts then carry the information to the parietal cortex and thereafter the dorsal horizontal tracts carry the information from the parietal cortex to the frontal cortex. The information likely carries onwards through the frontal cortex (FAT). According to the input-driven model of white matter development, the developmental progression of white matter tracts would be expected to proceed in this order: VOF →VHP →PVP →DHP →FAT.

The few studies that have looked at the development of white matter in the PVP have focused on the relationship between the microstructural tissue properties of the pArc and literacy. An association between FA, a common measure of tissue microstructure, in the pArc and reading is often reported in children 7 years and older (Deutsch et al., 2005; Huber et al., 2018; Klingberg et al., 2000; Wandell & Yeatman, 2013; Y. Wang et al., 2017; Yeatman, Dougherty, Ben-Shachar, et al., 2012; Yeatman et al., 2011). Research concerning behavioral correlates of PVP microstructure in children younger than 7 years, however, is extremely limited but suggests a similar association. One cross-sectional study in 5-8 year old children found that the microstructure of the pArc correlated positively with literacy (Broce et al., 2019). Another study using children of the same age range found that one year of literacy instruction resulted in microstructural changes in white matter voxels that likely corresponded to the PVP. In addition, the magnitude of the microstructural change correlated positively with children’s literacy gains over the same one-year period (Moulton et al., 2019). While evidence for a relationship between literacy and microstructure of tracts in the PVP in children younger than 7 years is still emerging, current work suggests that the information conveyed along the PVP is likely related to the early stages of learning to read.

Most prior work has focused almost exclusively on horizontal white matter tracts contained *within* regions associated with the ventral or dorsal streams (Bonekamp et al., 2007; Klaver et al., 2011; C. Lebel et al., 2008; Loenneker et al., 2011; Peters et al., 2014; Pierpaoli & Basser, 1996; Stiles et al., 2013; Yeatman, Dougherty, Myall, et al., 2012) the exception of the few studies reporting correlations between the pArc and behavior mentioned above (Deutsch et al., 2005; Huber et al., 2018; Klingberg et al., 2000; Wandell & Yeatman, 2013; Y. Wang et al., 2017; Yeatman, Dougherty, Ben-Shachar, et al., 2012; Yeatman et al., 2011). Consequently, we know very little about the development of PVP tracts that support the transfer of information *between* regions associated with the ventral and dorsal streams. Furthermore, prior work has employed a tract-of-interest approach, studying the developmental trajectory of individual tracts instead of the developmental relationships among tracts. Consequently, we know very little about the relative development of ventral, dorsal, and PVP white matter tracts. More specifically, we do not know how the development of the PVP may be related to the development of the ventral and dorsal streams. Our goal was to address: (1) the paucity of studies concerning the development of tracts within the PVP and (2) the developmental relationships among the PVP, ventral tracts, and dorsal tracts. Filling these gaps can provide insight into the mechanisms underlying white matter development as well as mechanisms of interaction between dorsal and ventral streams.

### Input-driven model of development in major white matter pathways

We propose an input-driven model of white matter development by which some portion of the underlying process of white matter development is driven by sensory input (**Figure 1b**). The model makes a step towards understanding the mechanisms that may underlie the development of white matter communication pathways connecting cortical regions with other cortical regions (i.e., cortico-cortical white matter tracts). The insight behind the model can be most simply explained as “input-driven” because it proposes that sensory input is one of the driving forces behind white matter development. According to the model, we would expect white matter serving processing stages that mature early (i.e., occipital cortex) to mature earlier than white matter serving processing stages that mature late (i.e., temporal and parietal cortices; **Figure 1b;** (Felleman & Van Essen, 1991; Hegdé & Felleman, 2007; Lerner et al., 2001; Meyer, 1981a, 1981b; Poggio & Ullman, 2013; Riesenhuber & Poggio, 1999; Serre et al., 2007)). Hence, tracts communicating within the occipital lobe (1 in **Figure 1b**) and carrying information out of the occipital lobe (2 in **Figure 1b**) will mature earlier than tracts carrying information out of the ventral cortex (3 in **Figure 1b**). Using the same reasoning, tracts sending information out of the ventral cortex will mature earlier than tracts sending information out of the parietal cortex (4 in **Figure 1b**) and tracts within the frontal cortex (5 in **Figure 1b**). The proposed model of white matter development is consistent with the understanding of the role of white matter in learning and plasticity that has shown that white matter microstructure changes with increased use and training (Bengtsson et al., 2005; Johansen-Berg et al., 2012; Sampaio-Baptista et al., 2013, 2018). The model is also consistent with aspects of the historical model of myelination and development proposed by Flechsig (Meyer, 1981a, 1981b).

Our model assumes that information from the occipital cortex travels to the ventral cortex and, from there, to the parietal cortex via the Posterior Vertical Pathway (PVP; **Figure 1b**). We note that the dorsal and ventral streams and their potential interactions are generally delineated based on functional and behavioral criteria (Culham et al., 2003; M. A. Goodale & Milner, 1992; T. W. James et al., 2003; Milner & Goodale, 2008; Mortimer Mishkin et al., 1983; Ungerleider & Haxby, 1994). Because of this, a major assumption in the literature has been that both the dorsal and ventral streams communicate *directly* from occipital to parietal and from occipital to ventral cortices, respectively (Binkofski & Buxbaum, 2013; Choi et al., 2020; Milner, 2017; Rizzolatti & Matelli, 2003). There is, however, currently very little evidence for major white matter tracts between the occipital and parietal cortices in humans. The lack of evidence for major tracts between the occipital and parietal cortices is in stark contrast to an abundant literature on the major tracts between occipital and ventral cortices, such as the Inferior Longitudinal Fasciculus (ILF; (Catani et al., 2002, 2003; Latini et al., 2017; Lawes et al., 2008; Mori et al., 2002, 2008; Ortibus et al., 2012; Panesar et al., 2018; Tusa & Ungerleider, 1985). Although it is possible that a series of local U-fibers may convey information between occipital and parietal cortices, as is the case in macaque (Baizer et al., 1991; Felleman & Van Essen, 1991; Majka et al., 2020), these U-fibers have not been characterized in humans and are likely to be pluri-synaptic. In sum, there has been no report of a major white matter tract directly connecting occipital and parietal cortices in over 100 years of inquiry (Catani et al., 2003; Choi et al., 2020; Jeremy D. Schmahmann, 2006; Kaneko et al., 2020; Takemura et al., 2015, 2019; Yeatman et al., 2014). One recent study specifically sought to find such a tract and was left wanting (Choi et al., 2020).

The input-driven model of white matter development proposed here (**Figure 1**) suggests that major white matter pathways develop in a particular order. The specific order that it proposes is that white matter tracts in occipital cortex (i.e., VOF) and in the ventral stream (i.e., Ventral Horizontal Pathway, VHP, including the ILF and IFOF) develop early, followed by tracts that connect the ventral stream to the dorsal stream (i.e., Posterior Vertical Pathway, PVP, including the pArc, TP-SPL, MDLFang, MDLFspl), and finally tracts in the dorsal stream (i.e., the Dorsal Horizontal Pathway, DHP, including the SLF1, SLF2, SLF3) and frontal cortex (FAT; **Figure 1a**). This developmental progression is consistent with two empirical findings. First, the microstructural tissue properties of white matter in the ventral-temporal cortex (specifically, the ILF) reaches adult-like levels in early childhood while the microstructural properties of dorsal white matter in parietal and frontal cortices (specifically, the SLF1, SLF2, SLF3) undergo more prolonged developmental trajectories (Klaver et al., 2011; C. Lebel et al., 2008; Loenneker et al., 2011; Stiles et al., 2013). Second, adult-like cortical function in posterior ventral-temporal cortices develops early compared to parietal and frontal cortices (Dekker et al., 2011; Golarai et al., 2007; Scherf et al., 2007); but see also (Karin H. James & Kersey, 2018)). Based on the known anatomical connections (see discussion in paragraph above), we would expect visual input to enter into the occipital cortex and then ventral-temporal cortex before proceeding onward to the parietal cortex and then frontal cortex, put explicitly: VOF →VHP →PVP →DHP →FAT (**Figure 1b**).

To fill gaps in knowledge about PVP development and to provide evidence for our input-driven model of white matter development (**Figure 1**), we collected diffusion-weighted magnetic resonance imaging (dMRI) data in 24 children (5-8 years old) and 13 adults (18-22 years old) to obtain an estimate of fractional anisotropy (FA) for each of the aforementioned tracts and pathways in children and in adults. We selected 5-8 years as the age range for our child sample because white matter microstructure and cortical function have been shown to be adult-like in the ventral-temporal cortex by 5 years of age (Klaver et al., 2011; C. Lebel et al., 2008; Loenneker et al., 2011; Stiles et al., 2013). The FA of the combined SLF1, 2 and 3, on the other hand, increases relatively slowly and plateaus at an adult-like measurement closer to 20 years of age (Klaver et al., 2011; C. Lebel et al., 2008; Loenneker et al., 2011; Stiles et al., 2013). Given that white matter microstructure in the VOF and VHP would be expected to be adult-like in the 5-8 year old age range, the input-driven model of white matter development proposed here (**Figure 1**) would predict that the next pathway to become adult-like would be the PVP. The model would further predict that the PVP should be more adult-like than the DHP and FAT white matter tracts in children 5-8 years of age.

## Materials and Methods

### Participants

Children and adult participants were recruited for this study. Children (5-8 yrs., n = 50) were recruited through an in-house database of parents in the local community who had expressed interest in participating in developmental psychological research and through word-of-mouth. Literate adult participants (18-25 yrs., n = 17) were recruited through an in-house database of IU students who had expressed interest in participating in psychological research at IU and through word-of-mouth. Of these, we obtained diffusion data from 31 children and 13 adults. Additional participant diffusion data were removed based on signal-to-noise (SNR) and/or motion concerns (see **Magnetic Resonance Imaging Data Analyses**), leaving 24 children (age: M = 6.7 years, SD = 1.3 years, Range = [4.7, 8.4], 15F, 9M) and 12 adults (age: M = 20.1 years, SD = 0.9 years, Range = [18.3, 21.2], 6F, 6M).

All participants were screened for neurological trauma, developmental disorders, and MRI contraindications. All participants were right-handed with English as their native language. Child participants were compensated with a small toy or gift card. Adult participants were compensated with a gift card.

### Behavioral Assessment

Children completed a behavioral session consisting of a battery of standard assessments designed to assess visual-motor integration (Beery VMI: green, blue, and brown; (Beery, 2004)), fine motor skill (Grooved Pegboard; (Matthews & Klove, 1964; Merker & Podell, 2011)), and literacy level (WJ-IV Achievement: Letter-Word Identification, Spelling, Word Attack, Spelling of Sounds; (Schrank & Wendling, 2018)). Assessments were administered in the same order for all participants. Raw scores for the Grooved Pegboard were measured in seconds to completion. All other raw scores were measured as the number of correct items.

A composite score quantified the abilities of each participant on three dimensions: visual-motor skill, fine motor skill, and literacy. The visual-motor composite score (VM) was calculated by averaging the percentage of correct responses on the Beery Visual-Motor Integration, Beery Visual Perception, and Beery Motor Coordination assessments. The fine motor skill composite score (FM) was calculated by averaging the time taken on the Grooved Pegboard for both hands, dividing by the number of rows completed (i.e., the children only complete two rows whereas the adults complete five rows), taking the inverse to make higher scores correspond to higher skill, and, finally, multiplying by one hundred to scale the score. The literacy composite score (LIT) was calculated by averaging the percentage of correct responses on the Woodcock Johnson IV Letter-Word Identification, WJ-IV Spelling, WJ-IV Word Attack, and WJ-IV Spelling of Sounds.

### Imaging Procedure

Child participants were first acclimated to the MRI environment by practice sessions in a full-sized and fully-equipped mock MRI simulator. The mock training session emphasized the importance of staying still during the scanning session. Children who completed the mock training session with success and comfort were then escorted into the MRI environment. Adult participants did not participate in the MRI simulator training session. All participants wore a Wheaton^®^ elastic shoulder immobilizer to reduce motion and wore ear protection. When space allowed, we used an inflatable head immobilization padding in the head coil. Participants were allowed to watch a movie, listen to an audio book, or to simply rest during scanning. Participants that successfully completed the neuroimaging session were asked to complete an additional behavioral session within one week of the neuroimaging session.

### Magnetic Resonance Image Acquisition

Neuroimaging was performed at the Indiana University Imaging Research Facility, housed within the Department of Psychological and Brain Sciences with a 3-Tesla Siemens Prisma whole-body MRI using a 32-channel head coil. High-resolution T1-weighted anatomical volumes were acquired using a Turbo-flash 3-D sequence: TI = 900 ms, TE = 2.7 ms, TR = 1800 ms, flip angle = 9°, with 160 sagittal slices of 1.0 mm thickness, a field of view of 256 x 256 mm, and an isometric voxel size of 1.0 mm^3^. Total acquisition time was 5 minutes and 12 seconds.

Diffusion data were collected using single-shot spin echo simultaneous multi-slice (SMS) EPI (transverse orientation, TE = 83.60 ms, TR = 3495 ms, flip angle = 90 degrees, isotropic 1.5 mm resolution; FOV = LR 210 mm x 192 mm x 138 mm; acquisition matrix MxP = 140 x 128. SMS acceleration factor = 4, interleaved). Data collected in the AP fold-over direction were collected at two diffusion gradient strengths, with 38 diffusion directions at b = 1,000 s/mm^2^ and 37 directions at b = 2,500 s/mm^2^. We then collected 10 diffusion images at b = 0 s/mm^2^ in the PA phase-encoding direction to use for distortion corrections. The total acquisition time was approximately 6 minutes.

### Magnetic Resonance Imaging Data Analyses

All analysis steps were performed using open and reproducible cloud services on the brainlife.io platform (Avesani et al., 2019) except for the statistical analyses (see below) that were performed in Matlab R2019b using customized code. All data and analysis services are freely available on brainlife.io (**Table 1**). The code for the statistical analyses is available https://github.com/svincibo/PVP-development.

**Table 1.** Data, description of analyses, and web-links to the open source code and open cloud services used in the creation of this dataset can be viewed in their entirety here: https://doi.org/10.25663/brainlife.pub.23.

| Application | Github repository | Open Service DOI | Git Branch |
| --- | --- | --- | --- |
| HCP AC-PC Alignment | https://github.com/brain-life/app-hcp-acpc-alignment | doi.org/10.25663/bl.app.99 | 1.4 |
| Freesurfer Segmentation | https://github.com/brainlife/app-freesurfer | doi.org/10.25663/bl.app.0 | 1.7 |
| Distortion and motion Correction | https://brainlife.io/app/5e6e72838a20890d8a8e96af | doi.org/10.25663/brainlife.app.287 | master |
| dMRI Preprocessing | https://github.com/brain-life/app-mrtrix3-preproc | doi.org/10.25663/bl.app.68 | 1.5 |
| Tractography | https://github.com/brain-life/app-mrtrix3-act | doi.org/10.25663/bl.app.101 | 1.3 |
| Tract Segmentation | https://github.com/brainlife/app-wmaSeg | doi.org/10.25663/brainlife.app.188 | 3.3 |
| Tract Cleaning | https://github.com/brainlife/app-removeTractOutliers | doi.org/10.25663/brainlife.app.195 | 1.3 |
| Tract Analysis Profiles | https://github.com/brain-life/app-tractanalysisprofiles | doi.org/10.25663/brainlife.app.185 | 1.8 |
| Tract Statistics | https://github.com/brainlife/app-tractographyQualityCheck | doi.org/10.25663/brainlife.app.189 | 1.2 |

#### Anatomical (T1w) processing

Anatomical images were aligned to the AC-PC plane with an affine transformation using HCP preprocessing pipeline (Glasser et al., 2013) as implemented in the HCP AC-PC Alignment App on brainlife.io (bl.app.99). Images from child participants were aligned using a pediatric atlas created from 5-8 year old children (Fonov et al., 2011); images from adult participants were aligned to the standard MNI152 adult template (Glasser et al., 2013). AC-PC aligned images were then segmented using the Freesurfer 6.0 (Fischl, 2012) as implemented in the Freesurfer App on brainlife.io (bl.app.0) to generate the cortical volume maps with labeled cortical regions according to the Destrieux 2009 atlas (Destrieux et al., 2010).

#### Diffusion (dMRI) processing

AP phase-encoded and PA phase-encoded images were combined using FSL Topup & Eddy with customized lambda values (brainlife.app.287). Susceptibility- and eddy current-induced distortions as well as inter-volume subject motion were also corrected in this step. All other diffusion preprocessing steps were then performed using the recommended MRtrix3 preprocessing steps (Ades-Aron et al., 2018) as implemented in the MRtrix3 Preprocess App on brainlife.io (bl.app.68). PCA denoising and Gibbs deringing procedures were performed first. The volumes were then corrected for bias field and rician noise. Finally, the preprocessed dMRI data and gradients were aligned to each participant’s ACPC-aligned anatomical image using boundary-based registration (BBR) in FSL (Greve & Fischl, 2009).

Diffusion data collected from either age group (i.e., child or adult) were removed from the sample if the Signal-to-Noise Ratio (SNR) was less than 15 or if the Framewise Displacement, a widely used measurement of head movement (Ciric et al., 2017; Power et al., 2012), was greater than 2 mm (see **Supplemental Materials** for additional information).

#### White matter measurements

The microstructural properties of white matter tissue were estimated in a voxel-wise fashion based on preprocessed dMRI data. We measured fractional anisotropy (FA) by fitting the diffusion tensor model on single-shell data (Basser et al., 1994; bl.app.68). FA is a summary measure of tissue microstructure that is related to white matter integrity and has been demonstrated to be reliable across a wide age range (Bonekamp et al., 2007).

#### Tractography

Probabilistic tractography (PT) was used to generate streamlines. We used constrained spherical deconvolution (CSD) to model the diffusion tensor for tracking (Tournier et al., 2007, 2012). Tracking with the CSD model fit was performed probabilistically, using the tractography procedures provided by MRtrix3 Anatomically-constrained Tractography (ACT; (Smith et al., 2012; Takemura et al., 2016) implemented in brainlife.io (bl.app.101). We note that the probabilistic tracking method has been quantitatively compared to Ensemble Tractography performed using MRtrix2 (Takemura et al., 2016) that was used in an earlier paper on similar white matter tracts (Bullock et al., 2019). We provide a series of supplemental panels to demonstrate that the quality of the final tract segmentation in the current paper is similar, if not higher, than that obtained by Bullock et al. (2019) (**Supplemental Figure 3**). We generated 2 million streamlines at L_max_ = 8 and maximum curvatures = 35 degrees, parameters that were optimized for our tractography needs. Streamlines that were shorter than 10 mm or longer than 200 mm were excluded. The tractogram was then segmented using a recently developed segmentation approach (D. Bullock et al., 2019) implemented in brailife.io (brainlife.app.188) that uses a method similar to white matter query language (WMQL; (Wassermann et al., 2013, 2016). We note that for some of the participants we had difficulties in addressing motion and imaging artifacts in the inferior frontal cortices; therefore, in some participants the anterior aspect of the IFOF was not included in our analyses. All the files containing the processed data utilized in this article can be accessed at https://doi.org/10.25663/brainlife.pub.23, participants excluded due to imaging artifacts were not included in the data release but can be made available upon request.

#### Cleaning

Streamlines that were more than 4 standard deviations away from the centroid of each tract and/or 4 standard deviations away from the relevant tract’s average streamline length were considered aberrant streamlines and were removed using the Remove Tract Outliers App on brainlife.io (brainlife.app.195).

#### Tract profiles

Tract-profiles were generated for each major tract (Yeatman, Dougherty, Myall, et al., 2012) as well as the additional PVP tracts (D. Bullock et al., 2019) using the Tract Analysis Profiles app on brainlife.io (brainlife.app.185). We first resampled each streamline in a particular tract into 200 equally-spaced nodes. At each node, we estimated the location of the tract’s ‘core’ by averaging the x, y, and z coordinates of each streamline at that node. We then estimated FA at each node of the core by averaging across streamlines within that node weighted by the distance of the streamline from the ‘core’ (**Figure 2b**). Averages for each tract were calculated by averaging across the central 160 nodes, excluding the first and last 20 nodes to avoid partial voluming effects from the cortex. Group averages for each tract were calculated by averaging the whole-tract average.

**Figure 2.**
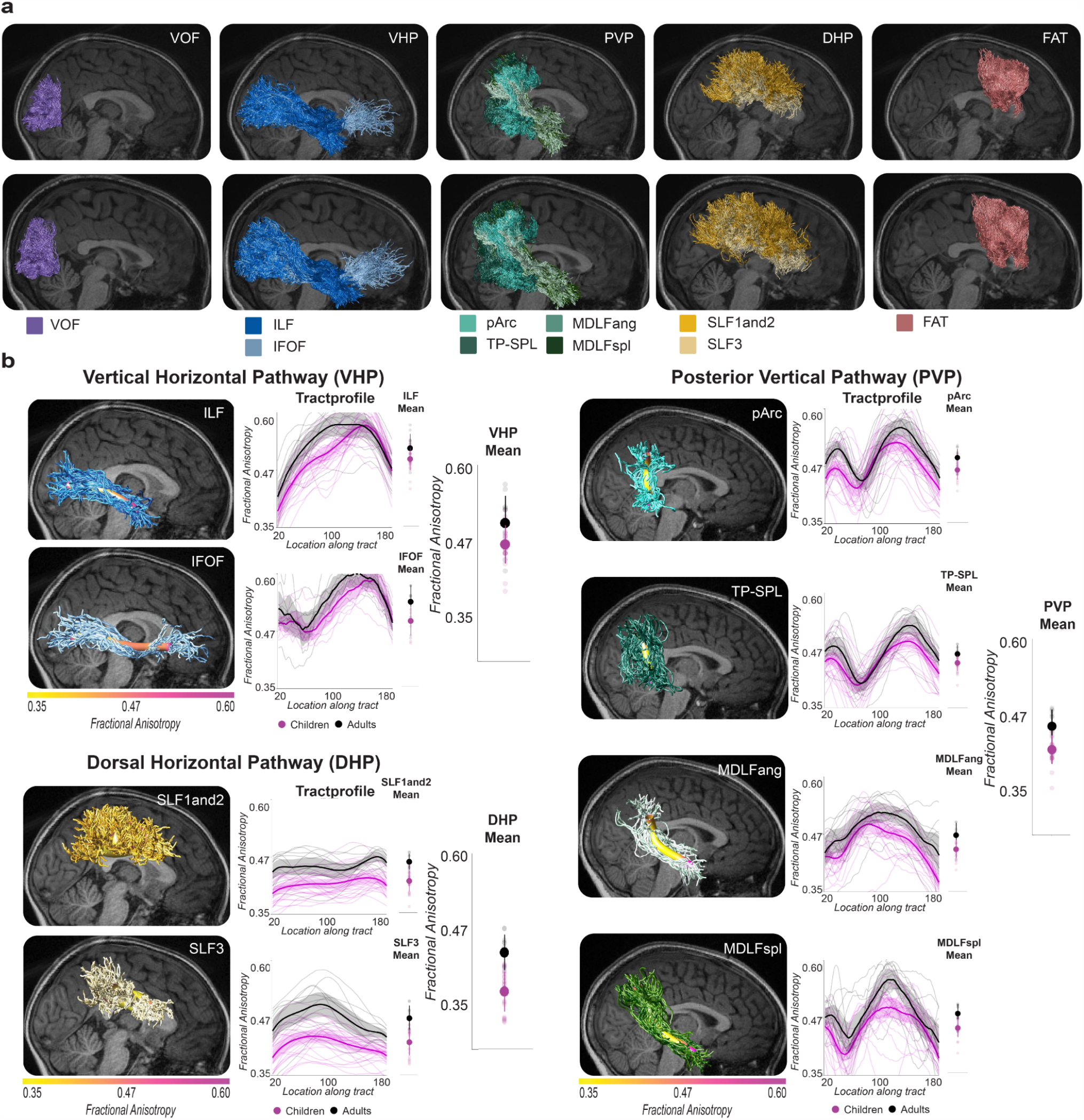
Description of Methods: White matter pathway definitions and white matter microstructure feature extraction. a. Pathways and tracts of interest. Pathways are displayed on representative child (top) and adult (bottom) brains. Both left and right hemispheres were evaluated; however, only the right hemisphere is displayed in the figure. Tracts were visualized using https://github.com/francopestilli/mba (Pestilli et al., 2014). **b. Tract profiles analysis**. FA is displayed on the y-axis; location along the tract ‘core’ is displayed on the x-axis. A tract ‘core’ was estimated for each tract in individual brains using the tractprofiles approach (Yeatman et al. 2012b) implemented as a service on brainlife.io (brainlife.app.185). To calculate the mean FA for a tract, we averaged across the core FA values, excluding the first and last 20 nodes to avoid partial voluming effects. To calculate the mean FA for a pathway, we averaged the mean tract FA values for tracts included in that pathway. For example, in the ventral horizontal (VHP) pathway, the mean FA for the ILF and IFOF were averaged to obtain the mean FA for the VHP pathway (see also **Figure 3**). We provide depictions of these analyses for each of the three pathways: the VHP pathway (top left), the dorsal horizontal (DHP) pathway (bottom left), and the posterior vertical pathway (PVP) (right). The solid line corresponds to the mean tract profile and the shaded areas correspond to 95% confidence intervals.

### Definition of Pathways

Individual white matter tracts were grouped into pathways to guide our analyses of the development of white matter within and between the ventral and dorsal streams. We focused on cortico-cortical white matter tracts. Tracts were assigned to pathways based on their anatomical location (posterior, anterior, dorsal, ventral) and cortical terminations (posterior, anterior, dorsal, ventral). Below we provide the rationale for how we assigned tracts to their pathways. We note that there are limitations in the spatial resolution of the diffusion image (Reveley et al., 2015; Thomas et al., 2014) that can limit the ability to capture fine details of the anatomy of the tracts. Yet, the tracts of interest here have been reported by diffusion imaging methods as well as tract-dissection methods and are largely consistent with comparative work (Kalyvas et al., 2020; Liu et al., 2020; Takemura et al., 2017).

#### Definition of horizontal white matter, ventral and dorsal pathways

We defined horizontal tracts as those tracts that connected anterior and posterior locations in the brain. These horizontal tracts included the IFOF, ILF, SLF1and2, and SLF3. The inferior longitudinal fasciculus (ILF) is a horizontal tract that connects the ventral temporal cortex with the occipital cortex (Catani et al., 2002; Nikos Makris et al., 2017). The inferior fronto-occipital fasciculus (IFOF) is a horizontal tract that connects the prefrontal cortex and occipital cortex (Lawes et al., 2008). The superior longitudinal fasciculus (SLF) is a large horizontal tract that connects the frontal motor and parietal cortex (de Schotten et al., 2011). The SLF can be segmented into two portions using available automated segmentation techniques (Bullock et al., 2019). SLF1and2 connects dorsal aspects of frontal cortex and superior parietal cortex; SLF3 connects ventral aspects of frontal cortex and superior parietal cortex.

Horizontal tracts were further separated into ventral (ILF, IFOF) or dorsal (SLF1and2, SLF3) tracts because prior work has demonstrated that dorsal tracts undergo a more prolonged developmental trajectory than ventral tracts (C. Lebel et al., 2008; Loenneker et al., 2011; Stiles et al., 2013). While ventral white matter is adult-like by about 5 years of age, dorsal white matter may not be adult-like until about 20 years of age (C. Lebel et al., 2008; Loenneker et al., 2011; Stiles et al., 2013). We, therefore, expected to find a developmental difference between ventral and dorsal horizontal tracts, such that the ventral tracts were more adult-like in our child sample then dorsal horizontal tracts. The Ventral Horizontal Pathway (VHP), therefore, contained the ILF and IFOF because these ventral tracts carry information between the posterior portion of the brain to the anterior portion (**Figure 2a**). The Dorsal Horizontal Pathway (DHP) contained the SLF1and2 and the SLF3 because these dorsal tracts carry information between the posterior portion of the brain to the anterior portion (**Figure 2a**).

#### Definition of vertical white matter, posterior vertical pathway and other vertical tracts

We defined vertical tracts as those tracts that connected dorsally and ventrally located cortical areas. These vertical tracts included the VOF, pArc, TP-SPL, MDLFspl, MDLFang, and FAT. The vertical occipital fasciculus (VOF) connects dorsal and ventral aspects of the occipital lobe (Rokem et al., 2017; Takemura et al., 2015; Weiner et al., 2017; Yeatman et al., 2014). The posterior arcuate (pArc) connects inferior parietal lobule with ventro-lateral portions of the middle portions of inferior and middle temporal gyri (Catani et al., 2005; Lawes et al., 2008; Weiner et al., 2017). The temporal parietal connection to the superior parietal lobe (TP-SPL) connects medial superior parietal lobule with ventro-lateral portions of the middle inferior temporal gyrus in the temporal cortex (Arash Kamali et al., 2014; Wu et al., 2016). The middle longitudinal fasciculus (MDLF) is a large vertical tract that connects the parietal cortex and anterior temporal cortex (A. Kamali et al., 2014). In humans, the MDLF has been reported to be separable into two subcomponents, connecting the anterior temporal lobe with the inferior and superior parietal lobule (N. Makris et al., 2013; Nikos Makris et al., 2017). MDLFspl connects superior parietal lobule with anterior temporal cortex; MdLFang connects the inferior parietal lobule with anterior temporal cortex (see (D. N. Bullock et al., 2021) for a more nuanced discussion). The frontal aslant tract (FAT) is an anterior vertical tract that connects posterior superior frontal gyrus and posterior inferior frontal gyrus (Dick et al., 2019; Lawes et al., 2008).

The vertical tracts were further separated based on the cortical regions that they connect. Two white matter tracts, the VOF and FAT, are tracts that connect cortical regions in the same lobe. The VOF connects cortical regions within the occipital lobe (Takemura et al., 2017) while the FAT connects cortical regions within the frontal lobe (Dick et al., 2019; Lawes et al., 2008). The pArc, TP-SPL, MDLFapl, and MDLFang all connect cortical regions in different lobes; they all connect cortical regions in temporal lobe to cortical regions in the parietal lobe. In other words, these four tracts directly connection cortical regions associated with the ventral and dorsal visual streams (D. Bullock et al., 2019; M. A. Goodale & Milner, 1992; M. Mishkin & Ungerleider, 1982). The Posterior Vertical Pathway (PVP), therefore, contained the pArc, TP-SPL, MDLFspl, and MDLFang because these tracts carry information between the ventral portion of the brain to the dorsal portion of the brain (**Figure 2a**). The PVP was of primary interest because tracts in this pathway directly connect cortical regions traditionally associated with the ventral and dorsal visual streams.

The VOF and FAT were selected as comparison tracts for two reasons. First, we wanted to ensure that our findings were specific to tracts connecting ventral and dorsal visual streams rather than some quality of vertical tracts more generally. Second, the VOF is in the posterior brain while the FAT is in the anterior brain and, therefore, these two tracts bookend the proposed developmental progression from early visual regions (i.e., VOF) to later regions (i.e., FAT; **Figure 1**).

### Experimental Design and Statistical Analyses

All statistical analyses were performed using either pathway group means or tract group means. To obtain a tract group mean, we first calculated a tract-average profile for each group, and then computed the mean across all nodes in the tract-average profile resulting in a single mean FA value for each tract and group. To obtain a pathway group mean, we calculated the average mean FA across tracts in each pathway, resulting in a mean FA value for each pathway in each age group (**Figure 2**).

All ANOVAs and multiple linear regressions were performed using SPSS Statistics for Mac OSX, version 25. All correlations and bootstrap tests were performed using Matlab R2019b. Matlab code that performs the post-processing analyses described in this study is available at https://github.com/svincibo/PVP-development. Additional details for each analysis are provided in the **Results** section.

#### Two-Way Repeated Measures ANOVA

We assessed differences in the developmental trajectory among white matter pathways (i.e., VOF, VHP, PVP, DHP, FAT) based on how adult-like these white matter pathways were in our child sample. We operationalized ‘adult-like’ as the difference in the average microstructural measurement of a pathway between children and adults, similar to other works (C. Lebel et al., 2008; Loenneker et al., 2011; Peters et al., 2014). A smaller difference would suggest that the pathway is more adult-like than a pathway with a larger difference between children and adults because the pathway with the larger difference score would be expected to be further from the adult-like measurement.

We first performed a 5 (Pathway: VOF, VHP, PVP, DHP, FAT) x 2 (Age Group: children, adults) Repeated Measures ANOVA to determine if there were any age group differences in white matter microstructure that depended on the pathway. The age-group difference was an estimate of how adult-like a particular pathway was in our sample. The smaller the difference, the more adult-like the microstructure of the pathway; the larger the difference, the less adult-like the microstructure of the pathway. The dependent variable was the mean FA along a tract’s profile, averaged across all tracts within that pathway and across hemispheres (**Figure 2**). The first factor, PATH, had five levels that were entered in this order: VOF, VHP, PVP, DHP, FAT. The second factor, AGE, had two levels: children and adults. PATH was within-participants and AGE was between-participants. SEX was entered as a covariate of no interest (Reynolds et al., 2019; Uda et al., 2015).

We predicted an interaction between AGE and PATH–that the magnitude of the difference in FA between children and adults would depend on the factor PATH. More specifically, we predicted a linear trend in the difference in FA between children and adults such that the size of the difference in FA between children and adults would follow the order proposed by the model (**Figure 1b**), namely, VOF, VHP, PVP, DHP, and FAT.

#### Multidimensional scaling and k-means clustering

Second, we performed multidimensional scaling and *k*-means clustering using a distance matrix calculated from the PVP-VHP and PVP-DHP correlations. Clustering is a method generally used to assess whether variables group together based on the similarity of some measure. Here, we assessed whether the PVP tracts grouped with the VHP tracts or the DHP tracts based on the similarity of their tissue microstructure. We tested the hypothesis that PVP microstructure was more similar to VHP microstructure than DHP microstructure in children by setting *k* = 2 and observing whether VHP or DHP tracts clustered with PVP tracts.

To prepare the data for *k*-means clustering, pairwise Pearson’s correlations (*r*) between tracts were computed based on z-scored FA values. The FA values were z-scored by computing the mean and standard deviation within each tract and group. We transformed the pairwise correlations into distance values (*d*) by computing the Euclidean distance between PVP-VHP and PVP-DHP correlation pairs and used *d* to perform classical multidimensional scaling (MDS; (Cox & Cox, 2008; Eisenberg et al., 2019; Kruskal, 1964; Seber, 2009; Torgerson, 1952).

MDS was used to decompose the distance values into independent dimensions that could be used for *k*-means clustering (Arthur & Vassilvitskii, 2006; Seber, 2009). A scree plot of the eigenvalues for each dimension in the MDS results and the final maximum relative error (*MRE*) of the model were used to select the appropriate number of dimensions (Cattell, 1966). Together, the scree plot and *MRE* values suggested that models using the first two dimensions were substantially better specified than those using only the first dimension in both data sets. Because of this, the first two dimensions were used for *k*-means clustering.

We used a silhouette coefficient to quantify the goodness of the *k*-means clustering solution. Silhouette scores range from -1 to 1. A silhouette score of 1 indicates that points in a cluster are densely grouped and very distant from points in other clusters. A silhouette score of 0 indicates that the points are not distinctly assigned to clusters; that the clustering solution is not significant, in other words. A silhouette score of -1 indicates that the points are likely assigned to incorrect clusters.

#### Behavioral correlations

To explore potential relationships between PVP microstructure and early literacy skills, a multiple linear regression analysis was performed to predict PVP microstructure in our child sample using three composite scores generated from different behavioral assessments that measured: literacy, visual-motor skill, and fine-motor skill (see **Behavioral Assessment** for a description of how the composite measures were computed). A large amount of work suggests an association between microstructure of PVP tracts and reading in children 7 years and older (Deutsch et al., 2005; Huber et al., 2018; Klingberg et al., 2000; Wandell & Yeatman, 2013; Y. Wang et al., 2017; Yeatman, Dougherty, Ben-Shachar, et al., 2012; Yeatman et al., 2011); however, our sample was 5-8 years of age. We, therefore, included visual-motor and fine-motor skill measures in the regression because a large amount of work has demonstrated that these two skills positively predict future literacy attainment (Cameron et al., 2012; Carlson et al., 2013; Clark, 2010; Dinehart, 2015; Fears & Lockman, 2018; Grissmer et al., 2010; Maldarelli et al., 2015). The dependent variable was the z-scored microstructural property (i.e., FA), averaged across tracts within the PVP, and the independent variables were the z-scored behavioral composite measures, as well as age in months and sex, excluding interaction terms. Only linear terms were included, because linear fits are most appropriate for this age range (Catherine Lebel et al., 2019). The regression model was tested for significance using an *F*-test of the overall model fit with the significance threshold set to *p* < 0.05. Each predictor was tested for significance using a one-sample *t*-test and a threshold of *p* < 0.05, using a Bonferroni correction for the 3 comparisons (i.e., the 3 predictors), *p*_*Bonferroni*_ = 0.05/3 = 0.017.

We performed two additional multiple linear regression analyses to determine, first, if any relationship between PVP microstructure and our composite scores was driven, first, by a particular tract within the PVP and, second, if and tract-specific relationship was driven by a particular subtest within the composite scores. Specifics of these follow-up regressions are provided with the reporting of the results.

## Results

### Vertical posterior pathway (PVP) tracts become adult-like before dorsal horizontal tracts

Our input-driven model of white matter development would predict that the development of white matter pathways should depend on the location of the pathway in the processing hierarchy, with pathways closer to sensory input (e.g., VOF or VHP) reaching adult-like microstructural values earlier than pathways further from sensory input (e.g, FAT or DHP; **Figure 1b**). More specifically, it predicts that in children 5-8 years of age, the PVP should be more adult-like than the DHP and the FAT.

We performed a Two-Way Repeated Measures ANOVA to determine if there were any age group differences in white matter microstructure that depended on the pathway. Results were in line with the prediction of the input-driven developmental model. FA in the PVP was more adult-like than in the DHP and FAT (**Figure 3**). This pattern of results suggests that the maturation of the pathways proceeds from early to late pathways in our proposed processing hierarchy. The Two-Way Repeated Measures ANOVA revealed an interaction between PATH and AGE, *F*(4, 132) = 4.109, *p* = 0.004. The main effect of PATH was significant, *F*(4, 132) = 19.473, *p* = 0.000. The main effect of AGE was significant, *F*(1, 33) = 26.928, *p* = 0.000. The significant interaction was driven by a linear trend, *F*(4, 33) = 9.723, *p* = 0.004; no other trends were significant. Post-hoc independent samples t-tests demonstrated that the significant linear trend was an increase in the difference in FA between children and adults across PATH, in the predicted order: VOF, *t*(34) = 3.013, *p =* 0.005, VHP, *t*(34) = 2.725, *p =* 0.010, PVP, *t*(34) = 4.297, *p* = 0.000, DHP, *t*(34) = 5.685, *p* = 0.000, and FAT, *t*(34) = 4.680, *p* = 0.000. All post-hoc independent samples t-tests passed Bonferroni correction for 5 comparisons at the *p* = 0.01 significance level (i.e., 0.05/5 = 0.01), except the *t*-tests for VOF and VHP tracts. PATH did not interact with SEX, *F*(4, 132) = 1.041, *p* = 0.389, our covariate of no interest. A Levene’s Test suggested that the error variance was not unequal across age groups, all *p*s > 0.05, indicating that a standard ANOVA and *t*-test were appropriate for these data.

**Figure 3.**
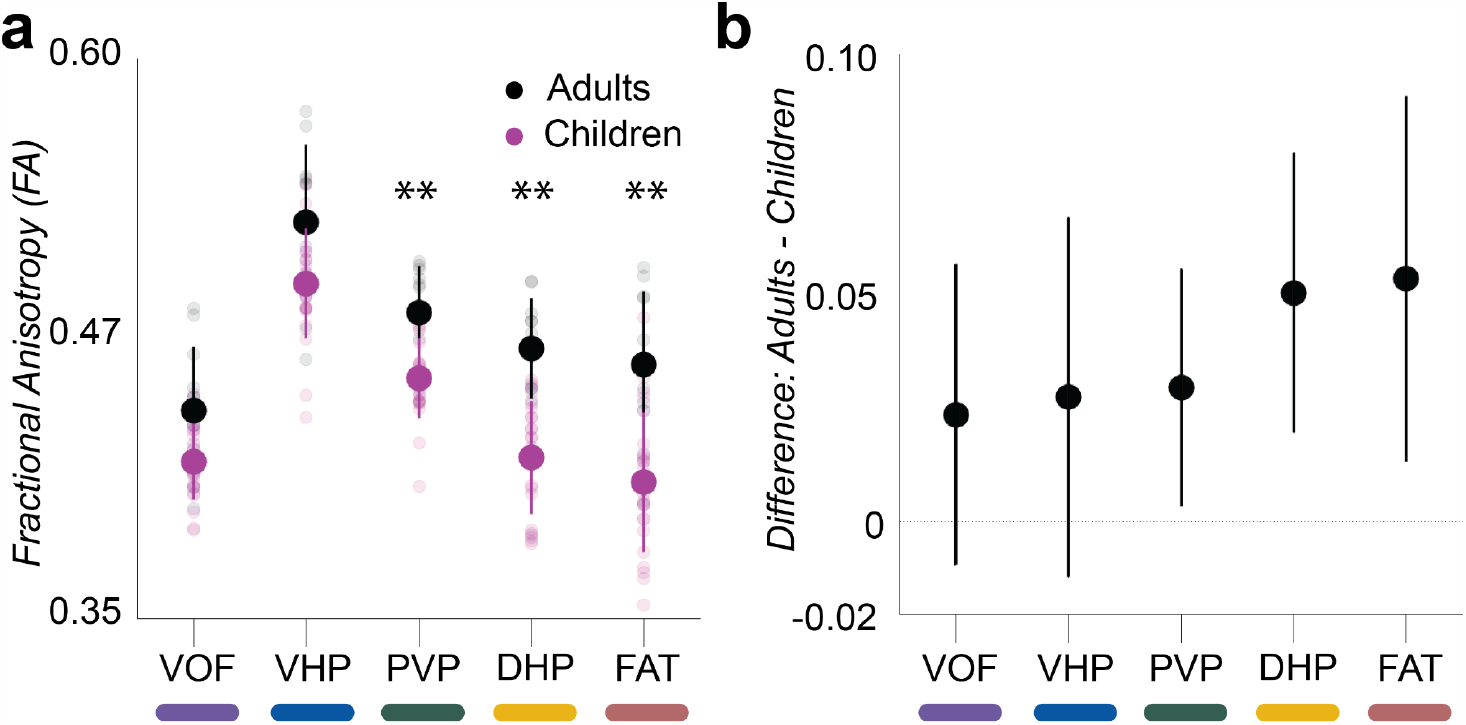
Interaction between Pathway and Age Group. **a. Group means for each pathway.** Group means for each pathway overlaid on individual participant means (transparent dots) color coded for age group. Adult means are shown in black and child means are shown in pink. The difference in fractional anisotropy (FA) between children and adults was dependent on Pathway. The age difference in FA increased linearly from the vertical-occipital fasciculus (VOF), to ventral horizontal tracts in the ventral visual stream (VHP), to the vertical posterior tracts that connect the ventral stream to parietal cortex (PVP), to dorsal horizontal tracts that connect parietal cortex to frontal motor cortex (DHP), and to the frontal aslant tract (FAT) that connects dorsal and ventral frontal motor cortex. **b. Difference in group means for each pathway**. Differences in group means are displayed for reference. Error bars represent standard deviation. **, *p* < 0.05 Bonferronni corrected.

### PVP tracts cluster with VHP tracts in children but not in adults

According to the input-driven model (**Figure 1**), during development the microstructure of a particular tract should be more associated with the tracts in the pathway from which it receives sensory inputs than tracts in pathways to which it sends inputs. The PVP was the first pathway in the proposed processing hierarchy to demonstrate a difference between children and adults in the previous analysis; it was preceded by an adult-like VHP and followed by a non-adult-like DHP (**Figure 3**). Based on our input-driven model of white matter development, we would expect that the microstructure of the PVP should be similar to the microstructure of the pathway that provides it with sensory input because simply carrying sensory input is expected to have a developmental effect in young children. More specifically, tracts carrying information out of the ventral cortex (PVP; 3 in **Figure 1b**) will be more similar to the mature tracts carrying information to the ventral cortex (VHP; 2 in **Figure 1b**) than the relatively immature tracts carrying information out of the parietal cortex (DHP; 4 in **Figure 1b**) in children.

To assess our prediction that PVP microstructure would be more similar to the microstructure of the upstream VHP relative to microstructure in the DHP in children, we applied a *k*-means clustering algorithm to the results of an MDS model of the correlations among tracts within the VHP, PVP, and DHP (**Figure 4a**). A scree plot of the eigenvalues for each dimension in the MDS result showed that both child and adult models were well specified by the first dimension (i.e., high eigenvalue in the first dimension relative to the other dimensions; **Figure 4b**). Including the second dimension, however, reduced the *MRE* by approximately half in both child and adult samples (first dimension alone: 0.67 in children and 0.55 in adults, first and second dimensions: 0.34 and 0.32; **Figure 4b**), indicating that including the second dimension produced a substantially better fit. We, therefore, applied the *k*-means clustering (*k* = 2) algorithm to the first two dimensions of the MDS model of the relations among tracts within the VHP, PVP, and DHP (**Figures 4a and 4b**).

**Figure 4.**
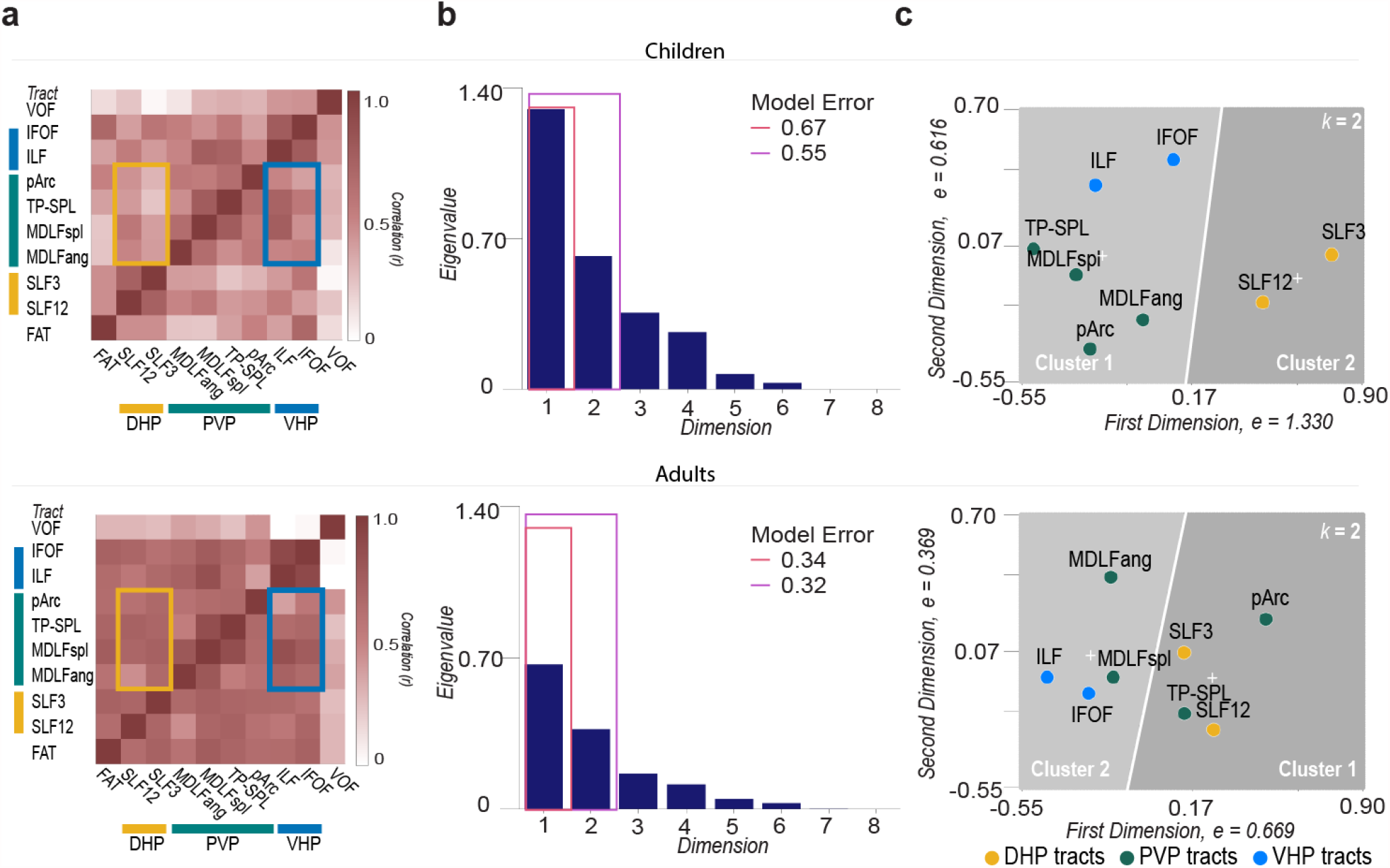
Multidimensional scaling of PVP-VHP and PVP-DHP correlations. **a. Pairwise correlations**. Pairwise correlations are between microstructure properties (FA) of tracts assigned to dorsal horizontal, ventral horizontal, and posterior vertical pathways. Tracts in the posterior vertical pathway (PVP) are indicated with a green bar. Tracts in the dorsal horizontal pathway (DHP) are indicated with a yellow box and bar and tracts in the ventral horizontal pathway (VHP) are indicated with a blue box and bar. In this analysis, a positive correlation between two tracts suggests that individuals with higher FA in one tract would also have higher FA in the second tract. Correlations among PVP and VHP tracts are framed in blue while correlations among PVP and DHP tracts are framed in yellow. All correlations were positive. **b. PVP correlations were modelled well using the first two MDS dimensions**. MDS eigenvalues were positive and large and suggested that the MDS models were able to explain a large portion of the variance in both child and adult data using only the first dimension. Eigenvalues were larger for the child data than for the adult data, suggesting that MDS was able to detect dimensionality better in the child data than in the adult data. The max relative error (*MRE*) values suggested that including the second dimension substantially increased the model fit. **c. Clustering of the MDS eigenvalues into two clusters**. *k*-means clustering with *k* = 2 was performed to test the hypothesis that PVP microstructure would be more similar to VHP than DHP microstructure. PVP tracts clustered with VHP tracts in the child data; however, this did not occur in the adult data (bottom). Cluster centroids are displayed as white crosses. Top panels refer to results in the children; bottom panels refer to results in the adults.

Results were as predicted. In children, PVP and VHP tracts clustered together while the DHP tracts clustered separately (**Figure 4c top**), suggesting that the microstructure of PVP tracts were more similar to the microstructure of VHP tracts than DHP tracts in children. The model *SSD* was 0.866. The PVP-VHP cluster had a silhouette score of 0.575 and the DHP cluster had a silhouette score of 0.834, indicating that the clustering solution was appropriate (see **Materials and Methods: Experimental Design and Statistical Analyses**). In adults, two PVP tracts clustered together with the VHP tracts, the MDLFspl and the MDLFang, and two PVP tracts clustered together with the DHP tracts, the pArc and the TP-SPL (**Figure 4c bottom**). The model *SSD* was 0.479. Cluster 1 had a silhouette score of 0.606 and cluster 2 had a silhouette score of 0.559.

The clustering solutions demonstrated that the PVP and VHP clustered together in children but not in adults (**Figure 4c**). To determine if the relationship among tracts within the VHP, PVP, and DHP pathways was statistically different in childhood compared to adulthood, we performed a bootstrap test on the PVP-VHP and PVP-DHP correlations to determine if the difference between VHP and DHP correlations in children was greater than the difference in adults, as suggested from the clustering solutions for children and adults. The result of this supplemental analysis was in line with what would be expected from the clustering solutions (**Figure 4c**). The difference between the PVP-VHP and PVP-DHP correlations in children was statistically greater than the difference in adults (**Supplemental Analysis 2.4** Bootstrap test of difference between children and adults, *z* = 1.841, *p* = 0.033). We note that this supplemental analysis cannot assess differences in the clustering solution because it does not test the clustering solution itself, a non-trivial pursuit (Amigó et al., 2009). Nonetheless, the results of the supplemental bootstrap testing support the conclusion that the relationship among tracts within the VHP, PVP, and DHP in childhood is different than the relationship in adulthood.

### Performance on a visual perceptual task predicts PVP microstructure in children

Results from Analyses 1 and 2 suggested that the age range of the child sample is likely a period of significant developmental change for the PVP. The PVP was the first white matter pathway in the proposed developmental progression (**Figure 1b**) that was not adult-like (**Figure 3**). The PVP microstructural properties were also more strongly associated with the microstructure of the VHP that precedes it than the DHP that follows it in the child sample (**Figure 4c**). These results and the input-driven model (**Figure 1**) suggest that PVP microstructural development might be driven by experience. Therefore, we investigated whether individual differences in PVP microstructure were related to behavioral measures of learning in the child sample. **Table 2** reports descriptive statistics for behavioral measures for children and adults.

**Table 2.** Descriptive Statistics and Behavioral Measures

|  | Age Group |  |
| --- | --- | --- |
|  | Children<br>(n=24) | Adults<br>(n = 13) |
|  | *M (SD)* | *M (SD)* |
| Age (years) | 6.7 (1.3) | 20.1 (0.9) |
| Beery |  |  |
| Visual-Motor Integration | 18.3 (3.6) | 27.1 (1.9) |
| Visual Perception | 20.9 (3.9) | 27.2 (2.6) |
| Motor Coordination | 18.1 (3.9) | 25.2 (2.6) |
| Grooved Pegboard |  |  |
| Dominant hand (right) | 42.4 (14.0)<sup>‡</sup> | 61.1 (8.5) |
| Non-dominant hand (left) | 45.7 (17.8)<sup>‡</sup> | 62.8 (6.2) |
| Woodcock Johnson IV |  |  |
| Letter Word Identification | 37.4 (21.0) | 70.5 (3.1) |
| Spelling | 17.3 (9.8) | 46.7 (4.3) |
| Word Attack | 15.8 (7.8) | 33.1 (21.8) |
| Spelling of Sounds | 11.1 (6.2) | 25.8 (2.9) |
| Composite Scores |  |  |
| Visual Motor | 19.1 (3.1) | 26.5 (1.3) |
| Fine Motor | 5.0 (1.5) | 3.3 (0.3) |
| Literacy | 20.4 (10.8) | 44.0 (6.5) |

*Behavioral testing occurred within 3 weeks of the neuroimaging session. Grooved Pegboard is reported in seconds to completion. All others are reported in a number of correct items. Composite scores were calculated by combining scores on more than one assessment, as reported in the main text.*

We first assessed the relationship between the average FA values across all tracts within the PVP and behavior. Results demonstrated that PVP microstructure was related to visual-motor and fine-motor skill in the child sample. FA in the PVP was significantly predicted by the model, *F*(5, 22) = 3.117, *p* = 0.035, with *R*^*2*^ = 0.478 and *R*_*adjusted*_ = 0.325. The full model and resulting parameters were: Predicted FA = 1.744 + 0.579(VM) - 0.420(FM) + 0.441(LIT) - 0.012(AGE) - 0.590(SEX). There were no significant predictors after Bonferroni correction for multiple comparisons, all *p*s > 0.017. AGE and SEX were also not significant predictors, all *p*s > 0.05.

To determine if a particular tract within the PVP was driving the relationship between PVP microstructure and the behavioral measures, a follow-up multiple linear regression was performed for each PVP tract separately, for a total of four regression models. Models were specified as described in the previous section and a Bonferroni correction was used to assess significance. Only one of the four models was significant. Results revealed a significant relationship between behavior and the pArc (**Figure 5a**), *F*(5, 22) = 4.435, *p* = 0.009, with *R*^*2*^ = 0.752 and *R*_*adjusted*_ = 0.438. The full model and resulting parameters were: Predicted FA = -0.217 + 0.782(VM) - 0.718(FM) + 0.109(LIT) - 0.011(AGE) + 0.794(SEX). Both VM and FM were significant predictors of FA in the pArc, VM: *t*(22) = 3.137, *p* = 0.006 and FM: *t*(22) = -2.822, *p* = 0.012. SEX, a covariate of no interest, was also a significant predictor of FA in the pArc, *p* = 0.016. We note that these results pass a multiple correction threshold correcting for the number of variables tested (3, *p* = 0.017), but not when correcting for the number of variables and models (4) tested altogether (12 total, *p* = 0.004). To conclude this model found a significant relationship between FA in the pArc and the visual-motor skill and fine-motor skill composite scores, although individual post-hoc tests did not pass multiple comparisons (**Figure 5a**).

**Figure 5.**
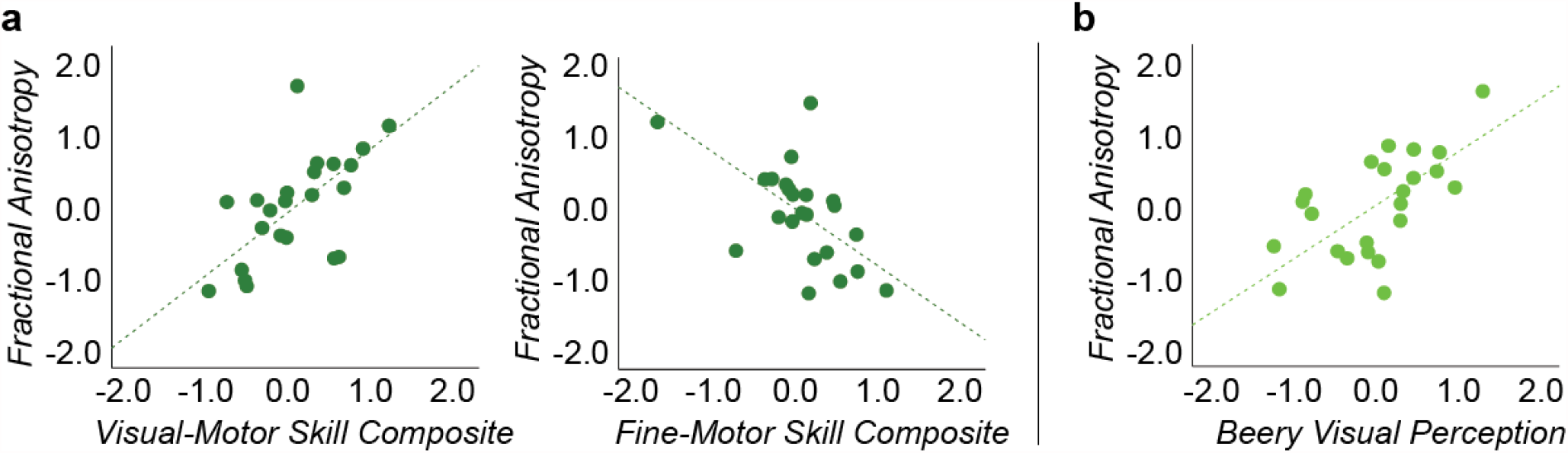
Performance on a visual perceptual task predicts FA in the posterior arcuate (pArc) in the child sample. **a. Relationship between pArc FA and Visual-motor and Fine-motor Skill**. Fractional anisotropy (FA) in the pArc was positively predicted by the VM composite score and negatively predicted by the FM composite score. **b. Relationship between pArc FA and performance on Beery Visual Perception**. Performance on the Beery Visual Perception (Beery VP) subtest positively predicted FA in the pArc. The Beery VP is one of three subtests used to calculate the VM composite score referenced in **a**. All measurements were based on the child sample and were z-scored.

To determine whether a particular subtest within the VM and/or FM composite scores was driving the relationship between these two composite measures and FA in the pArc, a follow-up multiple linear regression analysis was performed. The analysis included FA in the pArc as the dependent variable and 5 predictors (each predictor was a z-scored subtest used to calculate the original composite scores, see **Materials and Methods: Behavioral Assessment**). More specifically, the predictors consisted of the 3 subtests used to calculate the VM composite scores (Beery Visual Perception (VP), Beery Motor Coordination (MC), and Beery Visual-Motor Integration (VMI)) and the 2 subtests used to calculate the FM composite score (Grooved Pegs-R and Grooved Pegs-L). The LIT composite score was not tested because it was not identified as a significant predictor of FA in the pArc (see previous analysis). Age and sex were entered as covariates of no interest. A multiple comparison Bonferroni correction was applied (5 comparisons, one per predictor of interest, *p*_*Bonferroni*_ = 0.05/5 = 0.010). The model fit was significant, *F*(7, 22) = 3.410, *p* = 0.022, with *R*^*2*^ = 0.784 and *R*_*adjusted*_ = 0.614. The full model and resulting parameters were: Predicted FA = -1.1276 + 0.078(Beery VMI) + 0.644(Beery VP) + 0.135(Beery MC) + 0.577(GroovedPegs-R) - 0.259(GroovedPegs-L) + 0.005(AGE) + 0.104(SEX). Results revealed that performance on the Beery VP assessment significantly predicted FA in the pArc (**Figure 5b**; *t*(22) = 3.031, *p* = 0.008). GroovedPegs-R was marginally significant, but did not pass multiple comparisons corrections, *t*(22) = 2.171, *p* = 0.046. There were no other significant predictors, all *p*s > 0.010. We conclude that FA in the pArc of our child sample can be predicted by Beery Visual Perception scores; as FA increases, performance on Beery Visual Perception increases.

## Discussion

Our results, collectively, support an input-driven model of white matter development and suggest that the PVP might facilitate interactions between dorsal and ventral visual streams during learning and development (Freud et al., 2019; Hanisch et al., 2001). We first demonstrated that the difference between child and adult microstructure increased from early to late pathways: VOF, VHP, PVP, DHP, FAT. The PVP was the earliest pathway to demonstrate a difference in microstructure between children and adults, suggesting that the PVP was actively undergoing developmental change in our child sample. We then demonstrated that PVP microstructure was similar to VHP microstructure in our child sample. The results of this second analysis, considered alongside the developmental context established by the first analysis, suggest that the development of white matter pathways may be related to the communications received from relatively mature pathways. Finally, we demonstrated that FA in the pArc, a PVP tract connecting posterior ventral-temporal and inferior parietal cortices, was uniquely predicted by performance on a perceptual matching task, suggesting that the PVP may facilitate interactions between ventral and dorsal visual streams during learning.

### Vertical posterior pathway (PVP) tracts become adult-like before dorsal horizontal tracts

The development of horizontal tracts *within* the dorsal and ventral streams have been studied extensively (see reviews on gestation (Dubois et al., 2014; Huang & Vasung, 2014), infancy (Dubois et al., 2014; Qiu et al., 2015), childhood and adolescence (Catherine Lebel et al., 2019; Catherine Lebel & Deoni, 2018), adulthood (Assaf & Pasternak, 2008; S. Wang & Young, 2014), and geriatrics (Moseley, 2002)). A consistent finding from these prior works is that the microstructure of white matter in the ventral horizontal pathway (VHP) reaches adult-like levels in early childhood while the microstructure of the dorsal horizontal pathway (DHP) undergoes a more prolonged trajectory (C. Lebel et al., 2008; Loenneker et al., 2011; Stiles et al., 2013). Consistent with this prior literature, the current results revealed that the microstructure of tracts in the VHP was adult-like in our 5-8 year old child sample while the DHP was not yet adult-like. Thus, the current results and the prior literature support a ventral-to-dorsal developmental trajectory, where ventral white matter develops earlier than dorsal white matter (Klaver et al., 2011; C. Lebel et al., 2008; Loenneker et al., 2011; Stiles et al., 2013)..

To our knowledge, the current study is the first study to investigate the development of the recently described vertical tracts *between* the ventral and dorsal cortex (D. Bullock et al., 2019; Kalyvas et al., 2020; Arash Kamali et al., 2014; Nikos Makris et al., 2009; Maldonado et al., 2013; Menjot de Champfleur et al., 2013; Takemura et al., 2015; H. Wang & Yushkevich, 2013; Yeatman et al., 2014). We investigated the vertical-occipital fasciculus (VOF) connecting ventral and dorsal portions of the occipital lobe (Rokem et al., 2017; Takemura et al., 2015, 2017; Weiner et al., 2017; Yeatman et al., 2014) and also the frontal aslant tract (FAT) connecting ventral and dorsal portions of frontal motor cortex (Dick et al., 2019; Lawes et al., 2008). Our results revealed that the microstructure of the VOF was adult-like in our 5-8 year old child sample while the FAT was not yet adult-like. Furthermore, the posterior vertical pathway (PVP) containing four vertical tracts just anterior to the VOF was also not yet adult-like but was closer than the more anterior FAT. Thus, the current results support a posterior-to-anterior developmental trajectory.

Finding that the PVP was not yet adult-like in the 5-8 year old age range suggests that the PVP microstructure is undergoing active development in this age range. Our results demonstrated that the microstructure of the PVP was not adult-like in our child sample and we found a significant relationship between age and the microstructure of the PVP pathway in the child sample. As age increased, fractional anisotropy (FA) also increased (see **Supplemental Analysis 1.3**). However, there were two indications that the change in FA during childhood is moderate, which would suggest a more prolonged developmental trajectory. First, the slope of the regression line was small (0.37). Second, we did not find a significant difference in FA between younger children (4.5-6.5 years) and older children (6.5-8.5 years). Thus, it is likely that the PVP undergoes a more prolonged developmental trajectory with significant changes happening during adolescence, similar to the DHP white matter.

Observing the development of horizontal tracts and the development of vertical tracts together adds context to the development of individual tracts. The ventral-to-dorsal developmental trajectory of horizontal tracts interweaves with the posterior-to-anterior developmental trajectory of the vertical tracts to suggest a more nuanced understanding of white matter development. More specifically, our results suggest that cortico-cortical white matter develops along a ventral/posterior-to-dorsal/anterior trajectory which is a developmental pattern where the white matter tracts closer to sensory inputs develop earlier than tracts further from sensory input. This developmental pattern is exactly the pattern that would be expected if some portion of white matter development were driven by the sensory input (in this case visual input). We, therefore, interpret our findings to suggest that the microstructural development of white matter pathways is in some way related to the propagation of sensory information.

### PVP tracts cluster with VHP tracts in children but not in adults

To better understand the developmental relationship between the PVP and nearby white matter pathways, we focused on the PVP and the pathways immediately upstream (VHP) and downstream (DHP) of the PVP. Based on our model of white matter development, we hypothesized that development of the PVP was related to the incoming data from cortical regions receiving input from the relatively mature VHP. Therefore, we predicted that the microstructure of the PVP would be more similar to the VHP that precedes it than to the DHP that follows. Because this prediction was related to how the white matter develops, we expected to find this in the child sample, but not in the adult sample. Results were in line with our predictions. The PVP tracts clustered with the VHP tracts in the child sample but not in the adult sample, suggesting that tracts within the VHP, PVP, and DHP pathways have a different relationship in childhood than in adulthood.

Our results demonstrate that the microstructure of white matter pathways connecting dorsal and ventral streams is related to the microstructure of white matter pathways in the ventral stream in 5-8 year old children. Our interpretation of this result is that tracts within the VHP and PVP are microstructurally similar, indicating that they might have similar information-transfer properties (e.g., timing and bandwidth of information transfer might be similar; (Drakesmith et al., 2019)). The cortical region that unites the VHP and PVP pathways is ventral-temporal cortex; thus, our interpretation of these results is that the VHP and PVP pathways are both important for carrying signals among occipital, ventral-temporal, and parietal cortices that are related to the development of function in ventral-temporal cortex. Indeed, an emerging line of research suggests that anatomical connectivity predicts the location of category selective responses in the ventral-temporal cortex (Li et al., 2020; Osher et al., 2016; Saygin et al., 2011, 2016). For example, one such study found that the anatomical connectivity of the visual word form area (VWFA) at age 5 predicted the functional location of the VWFA at age 8 (Saygin et al., 2016). In the context of these prior works, our results add that a similarity in microstructure of anatomical connections with the ventral-temporal cortex may be important for coordinating signals in a way that supports the development of category selectivity in the ventral-temporal cortex in the 5-8 year old age range.

The relationships among the VHP, PVP, and DHP tracts were different in adulthood than they were in childhood. In adulthood, two of the PVP tracts clustered with the VHP tracts (similar to childhood) and two clustered with the DHP tracts (different from childhood). The two tracts that clustered with the DHP in adulthood were the TP-SPL and pArc, both of which terminate in the posterior ventral-temporal cortex. In line with our interpretation of the clustering results in childhood, we propose that these results indicate that the TP-SPL and pArc might have similar communication transfer properties as the DHP in adulthood. Further research will be necessary to relate the information-transfer properties of these tracts (e.g., bandwidth and timing) by directly relating electrophysiological measurements and measurements of white matter properties.

### Performance on a visual perceptual task predicts PVP microstructure in children

Our final analysis was designed to characterize the relationship between the PVP and literacy. The purpose of this analysis was, first, to demonstrate that PVP development is relatable to behavior and, second, to contribute to our understanding of the nature of information communicated along PVP tracts. We found that FA in the posterior arcuate (pArc) that connects posterior ventral temporal cortex and inferior parietal lobe was predicted by children’s visual-motor integration composite scores and, more specifically, was predicted by performance on the visual perception subtask. We found no other behavioral associations with any other white matter pathway tested, suggesting some degree of specificity to the pArc and visual-motor skill. Our results suggest that the anatomical connectivity between the posterior ventral-temporal cortex and inferior parietal lobe may be related to the development of the visual perceptual aspect of visual-motor integration.

Many studies have found a relationship between the pArc and literacy in children 7 years and older (Deutsch et al., 2005; Huber et al., 2018; Klingberg et al., 2000; Wandell & Yeatman, 2013; Y. Wang et al., 2017; Yeatman, Dougherty, Ben-Shachar, et al., 2012; Yeatman et al., 2011). We did not find a similar association in our child sample. Instead, we found an association between the left pArc and the perceptual aspect of visual-motor skill. We attribute the apparent discrepancy between our results and the majority of the prior work to a difference in the age range of the child samples used. While our age range was 5-8 years, all but one prior work used children 7 years and older. We note that a breadth of behavioral work has indicated that visual-motor skill development positively predicts literacy development in this age range (Cameron et al., 2012; Carlson et al., 2013; Clark, 2010; Dinehart, 2015; Fears & Lockman, 2018; Grissmer et al., 2010; Maldarelli et al., 2015), suggesting that the left pArc’s relationship with visual-motor skill may be a precursor to its relationship with literacy. Nonetheless, the one prior work that used the same age range as our study also reported a relationship between pArc and literacy (Broce et al., 2019), suggesting that future work will be necessary to better understand the relationship between the pArc and literacy development (Yeatman & White, 2021).

## General Discussion

A significant amount of research in developmental neuroimaging focuses on the 5-8 year old age range as an important period for the development of adult-like category selectivity in ventral-temporal cortex (Cantlon et al., 2011; Cohen et al., 2019; Grill-Spector et al., 2008; Karin H. James, 2017; Longcamp et al., 2008; Saygin et al., 2016). Motor learning experiences are particularly important for the development of category selectivity in this age range ((Karin H. James, 2017; Longcamp et al., 2008; Wakefield & James, 2011); although category selectivity can emerge in the absence of motor learning (Striem-Amit et al., 2017)). Handwriting experience, for example, increases letter-selective activation in the ventral-temporal cortex in 5 year old children (Karin Harman James, 2010; Karin H. James & Engelhardt, 2012) and, at the same time, increases functional communication among the ventral-temporal, parietal, and frontal cortices (Vinci-Booher et al., 2016). Thus, it has been proposed that action increases object recognition abilities by integrating the perceptual system in ventral-temporal cortex with the motor system in parietal and frontal cortices (Karin H. James & Gauthier, 2006; Longcamp et al., 2008; Seger & Miller, 2010; Vinci-Booher et al., 2016).

A major gap in research on the role of action experience in the development of object selectivity has been the inability to ground the functional networks observed during object perception onto underlying anatomical pathways. Although the current work did not directly assess the relationship between function and anatomy during object recognition, the finding that the PVP is still developing in the 5-8 year old age range is consistent with the notion that the PVP supports communications among ventral-temporal and parietal cortices during object recognition. Thus, the current work suggests that functional communication between these two streams may be mediated by the white matter tracts connecting the cortical areas in each stream and may be important for cortical development in both streams. Understanding the role of white matter connections within and between the streams can help us understand their degree of interaction and how this might change over the course of the development.

Our model proposes that *some* portion of the white matter development should be related to the propagation of incoming sensory information, such that white matter closer to sensory input would be expected to develop earlier than white matter further from sensory input. This would be expected simply because white matter nearer to sensory input receives sensory signals more directly and more often than white matter further from sensory input. This interpretation is consistent with a large body of work that has demonstrated that sensory input is a major driver of developmental change in the brain (Flechsig Of Leipsic, 1901; Maurer & Lewis, 2018; Meyer, 1981a, 1981b): white matter develops earlier in tracts closer to sensory input and later in tracts further away from sensory input. We propose that this developmental trajectory is related, at least in part, to the propagation of sensory information along the visual hierarchy and the major white matter pathways subserving this hierarchy. However, it is important to note that diffusion imaging is limited in that it cannot provide information concerning the direction that information-flow along the tracts. Feedback communications from the parietal to the ventral-temporal cortex are also likely important for neural development, as discussed above for the case of the development of functional selectivity for object categories.

We note that this is the first attempt that we know of to understand the relationship among white matter tracts. While we have a general understanding of the developmental trajectories of different white matter tracts, our understanding of how these developmental trajectories interact over the course of development is limited. Such an understanding can provide insight into how the brain develops in relation to behavior. For example, the finding that the microstructure of major white matter pathways that support communications with ventral-temporal cortex is adult-like by the age of 5 years can help us understand how and why the emergence of category selectivity might occur between the ages of 5-8 years of age. More work is necessary to understand how the relationships among communication pathways support the relationship between brain and behavior over the course of development.

## Acknowledgments

This research was funded by NSF OAC-1916518, NSF IIS-1912270, NSF IIS-1636893, NSF BCS-1734853, Microsoft Investigator Fellowship to F.P. Data collections as supported by The Emergent Areas or Research Indiana University to F. Pestilli, K. James and L. Smith, the Indiana University Bloomington Imaging Research Facility Brain Scan Credit Program, the Indiana Clinical and Translational Sciences Institute, and the Johnson Center for Innovation and Translational Research provided additional imaging funds. SVB was partially supported by the NIH and institute T32-HD007475-21 and the EAR Initiative Linda Smith (Indiana University). DNB was supported by NIH NIMH T32-MH103213 to William Hetrick (Indiana University). The authors would like to acknowledge the help of Dr. Hu Cheng for help with the imaging sequences, Soichi Hayashi and Brent McPherson for contributing to the development of brainlife.io.

