## Supplemental Materials for "Development of white matter tracts between and within the dorsal and ventral streams"

**Abbreviated title:** White matter development between streams

**Authors' Names:** Vinci-Booher, S.*[1], Caron, B.[1], Bullock, D.[1], James, K.[1], & Pestilli, F.*[1,2]

**Author Affiliations:** [1]Indiana University, Bloomington, Indiana, USA; [2]University of Texas, Austin, Texas, USA

***Corresponding Authors:**

Franco Pestilli
Department of Psychology
The University of Texas at Austin
108 E Dean Keeton St, Austin, TX 78712
(347) 829- 4185,

Sophia Vinci-Booher
Department of Psychological and Brain Sciences
Indiana University
1101 E. 10th Street, Bloomington, IN 47405
(812) 855-1494,

**Acknowledgments**

This research was funded by NSF OAC-1916518, NSF IIS-1912270, NSF IIS-1636893, NSF BCS-1734853, Microsoft Faculty Fellowship to F.P. Data collections as supported by The Emergent Areas or Research Indiana University to F. Pestilli, K. James and L. Smith, the Indiana University Bloomington Imaging Research Facility Brain Scan Credit Program, the Indiana Clinical and Translational Sciences Institute, and the Johnson Center for Innovation and Translational Research provided additional imaging funds. SVB was partially supported by the NIH and institute T32-HD007475-21 and the EAR Initiative Linda Smith (Indiana University). DNB was partially supported by NIH NIMH T32-MH103213 to William Hetrick (Indiana University). The authors would like to acknowledge the help of Dr. Hu Cheng for help with the imaging sequences, Soichi Hayashi and Brent McPherson for contributing to the development of brainlife.io.

**Author Contribution Statement**

Sophia Vinci-Booher contributed to all aspects of the manuscript, including the original conception of the study, ongoing conceptual development, the design, data collection, analyses and software, writing the original draft of the paper, and revisions. Brad Caron and Dan Bullock contributed software and supported software development, participated in data quality checks, and commented on the manuscript. Karin James contributed to data collection, and commented on the manuscript. Franco Pestilli contributed to the original conception of the study, the conceptual development of the work, the design, the analyses, software, training of Sophia Vinci-Booher, and in the writing of the manuscript and revisions.

**Conflict of Interest Statement**

The authors declare no competing financial interests.

### Data processing and quality assurance

We performed a series of quality assurance steps to demonstrate how our approach to data preprocessing generates good quality features that can be used for the study we presented.

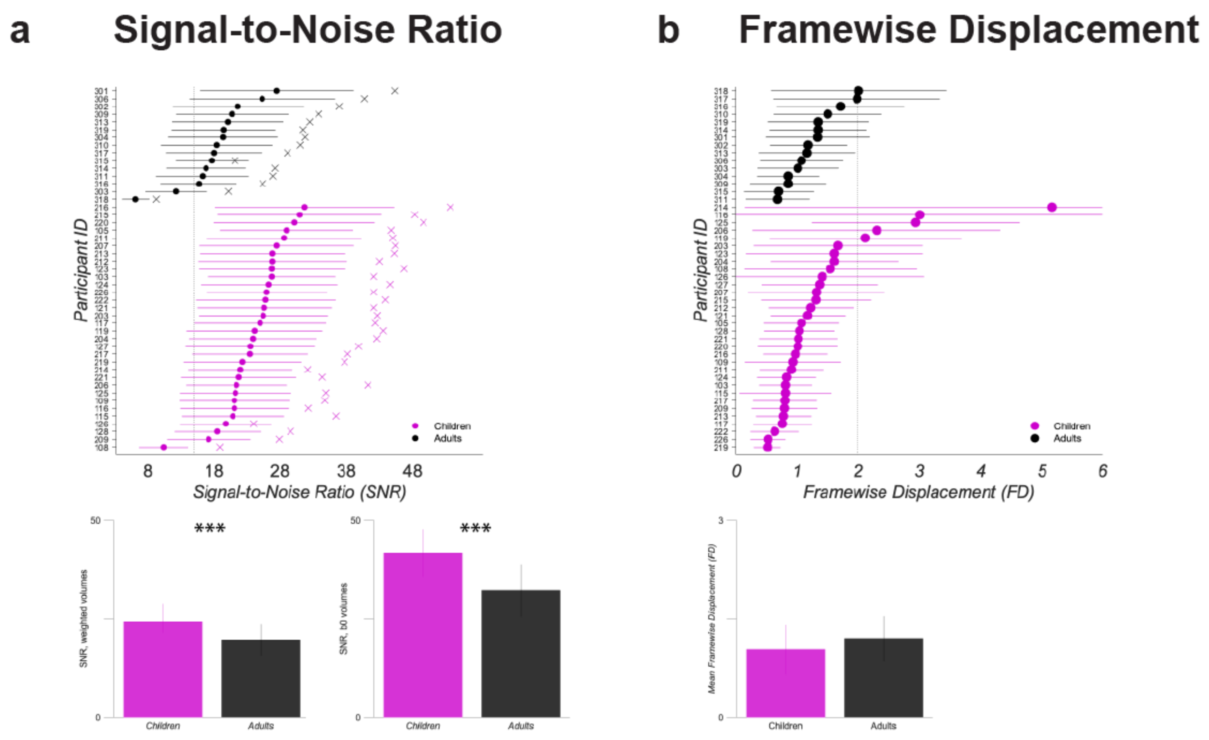


**Supplemental Figure 1. Quality Assurance of Diffusion Volumes. (a) Signal-to-Noise Ratio.** Signal-to-noise ratio (SNR) in the corpus callosum was calculated for each diffusion image to estimate image quality (https://brainlife.io/app/5bca308550fdf50028c6342c). Mean SNR of b0 volumes are indicated with an x and mean SNR of weighted volumes are indicated with a filled dot. Participants with images with SNR less than 15 (indicated by the vertical line) were excluded. This included 108, 303, and 318. We directly compared both mean SNR values between adults and children, including only participants whose data were analyzed in the main analyses (see main text). SNR of b0 volumes was significantly different between children and adults, $t(34) = 4.305$, $p = 0.0001$. SNR of b0 volumes was significantly different between children and adults, $t(34) = 4.104$, $p = 0.0002$. In both cases, SNR was greater in children than in adults. **(b) Framewise Displacement.** Framewise Displacement (FD) was calculated for each diffusion image to estimate head motion (https://brainlife.io/app/58c56cf7e13a50849b258800). Participants with images with FD greater than 2 (indicated by the vertical line) were excluded. This included 116, 119, 125, 206, 214, 317, and 318. We directly compared both FD values between adults and children, including only participants whose data were analyzed in the main analyses (see main text). FD was not significantly different between children and adults, $t(34) = 1.275$, $p = 0.2108$. We, therefore, did not include head movement estimates as covariates in our analyses. ***, $p < .001$.

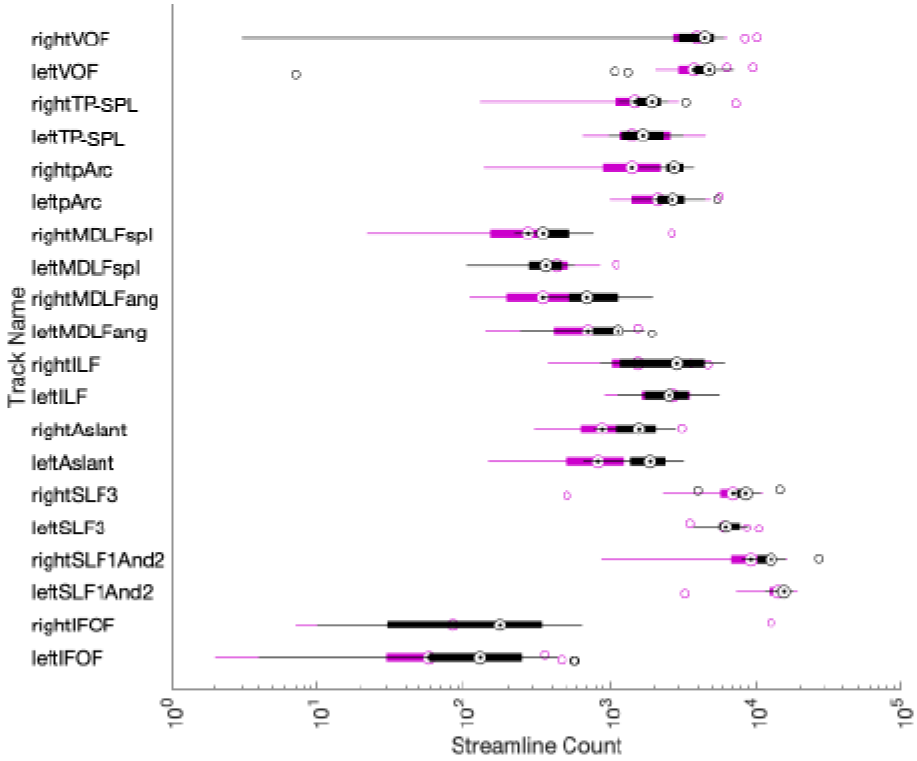


**Supplemental Figure 2. Quality Assurance of Tractography.** Box and whisker plots of streamline count for all tracts of interest are plotted for children (pink) and adults (black).

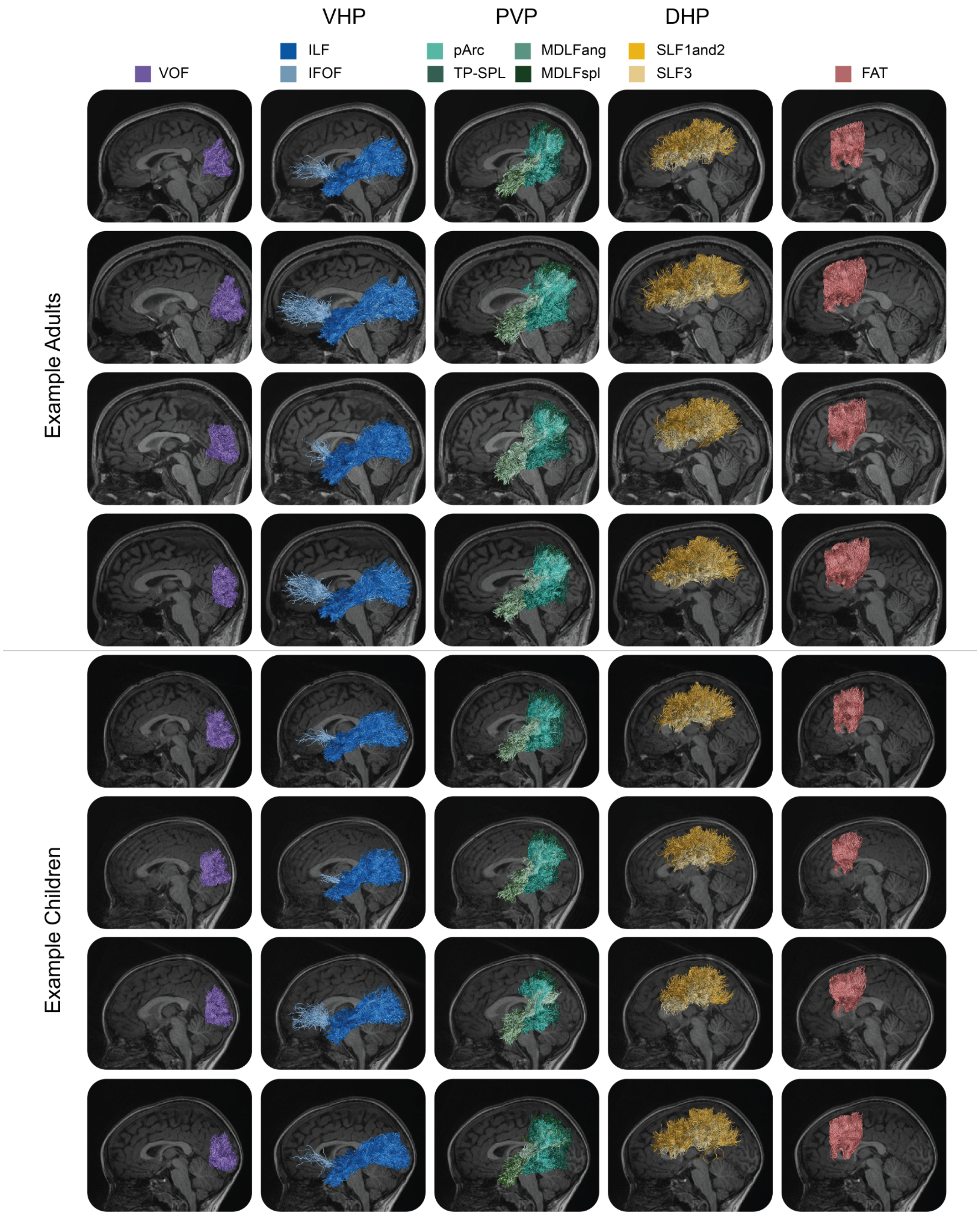


**Supplemental Figure 3. Tractography from example participants.**

**Supplemental Analyses for "Vertical posterior tracts become adult-like before dorsal horizontal tracts"**

Three analyses were performed to verify that the results reported in the main text did not depend on hemisphere (**Supplemental Analysis 1.1**), tract within pathway (**Supplemental Analysis 1.2**), and age within the child group (**Supplemental Analysis 1.3**). We found no evidence of hemispheric asymmetry, validating averaging across hemispheres. We found no evidence of tract-related heterogeneity within any pathway, validating averaging across tracts within a pathway. We found no evidence of age-related heterogeneity in our child sample, validating the combination of all children into one age group.

**Supplemental Analysis 1.1 Hemispheric differences**

We found no evidence of hemispheric asymmetry. A 5 (PATH: VOF, VHP, PVP, DHP, FAT) x 2 (Age Group: Children, Adults) x 2 (Hemisphere: Left, Right) Repeated Measures ANOVA with Sex entered as a covariate of no interest revealed a significant main effect for PATH, $F(4, 132) = 20.074$, $p = 0.000$, and a significant two-way interaction between PATH and Age Group, $F(4, 132) = 4.063$, $p = 0.004$, as in the original Two-Way Repeated Measures ANOVA. The three-way interaction, however, was not significant, $F(4, 132) = 0.396$, $p = 0.811$, post hoc observed power = 0.139, suggesting that the difference between children and adults among tract categories did not depend on hemisphere. Post hoc 5 (PATH: VOF, VHP, PVP, DHP, FAT) x 2 (Age Group: Children, Adults) Repeated Measures ANOVAs with Sex entered as a covariate of no interest were conducted for each hemisphere separately. These post hoc ANOVAs revealed a significant linear interaction in both the left, $F(1, 33) = 6.380$, $p = 0.017$, and the right hemispheres, $F(1, 33) = 9.116$, $p = 0.005$; both linear interactions were in the same direction, with the difference in FA between children and adults across PATH increasing in the predicted order: VOF, VHP, PVP, DHP, FAT.

**Supplemental Analysis 1.2 Pathway cohesion**

We found no evidence of tract-related heterogeneity within any pathway. We performed a Two-Way Repeated Measures ANOVA for each PATH that included two factors, Tract and Age Group. Tract had as many levels as tracts that were included in that PATH. Age Group always had 2 levels, children and adults. The dependent variable was always fractional anisotropy (FA). Results for each pathway are reported separately.

*Ventral Horizontal Pathway (VHP).* A Two-Way Repeated Measures ANOVA for Tract (L-ILF, L-IFOF, R-ILF, R-IFOF) and Age Group (children, adults) with Sex entered as a covariate of no interest revealed a significant main effect of Age Group, $F(1, 10) = 21.709$, $p = 0.001$. The main effect of Tract was not significant, $F(3, 30) = 0.743$, $p = 0.535$. The interaction between Tract and Age Group was not significant, $F(3, 30) = 1.207$, $p = 0.324$, post hoc power for alpha = 0.05 at 0.290, suggesting that the difference between children and adults did not differ among tracts that were included in the VHP.

*Vertical Posterior Pathway (PVP).* A Two-Way Repeated Measures ANOVA for Tract (L-pArc, L-TP-SPL, L-MDLFspl, L-MDLFang, R-pArc, R-TP-SPL, R-MDLFspl, R-MDLFang) and Age Group (children, adults) with Sex entered as a covariate of no interest revealed a significant main effect of Tract, $F(7, 224) = 3.913$, $p = 0.000$, and of Age Group, $F(1, 32) = 18.194$, $p = 0.000$. The interaction between Tract and Age Group was not significant, $F(7, 224) = 1.622$, $p = 0.130$, post hoc power for alpha = 0.05 at 0.665, suggesting that the difference between children and adults did not differ among tracts that were included in the PVP.

*Dorsal Horizontal Pathway (DHP).* A Two-Way Repeated Measures ANOVA for Tract (L-SLF1and2, L-SLF3, R-SLF1and2, R-SLF3) and Age Group (children, adults) with Sex entered as a covariate of no interest revealed a significant main effect of Age Group, $F(1, 33) = 30.584$, $p = 0.000$. The main effect of Tract was not significant, $F(3, 99) = 1.885$, $p = 0.137$. The interaction between Tract and Age Group was not significant, $F(3, 99) = 1.500$, $p = 0.219$, post hoc power for alpha = 0.05 at 0.386, suggesting that the difference between children and adults did not differ among tracts that were included in the DHP pathway.

**Supplemental Analysis 1.3 Age group cohesion**

We found no evidence of age-related heterogeneity in our child sample, validating the combination of all children into one age group: children. A 5 (PATH: VOF, VHP, PVP, DHP, FAT) x 2 (Age Group: Children 4.5 - 6.5 years, Children 6.5 - 8.5 years) Repeated-Measures ANOVA with Sex entered as a covariate of no interest revealed a significant main effect of PATH, $F(4, 84) = 13.250$, $p = 0.000$; however, neither the main effect of Age Group nor the interaction between PATH and Age Group were significant, $F(1, 21) = 2.878$, $p = 0.105$, post hoc power for alpha = 0.05 at 0.367, and $F(1, 21) = 0.399$, $p = 0.809$, post hoc power for alpha = 0.05 at 0.138, respectively. The covariate SEX was also not a significant predictor of FA, $F(1, 21) = .363$, $p = 0.553$, and SEX did not interact with PATH, $F(1, 21) = .011$, $p = 0.918$. The lack of a statistically significant difference between the two age-groups (Age Group: Children 4.5 - 6.5 years, Children 6.5 - 8.5 years) in the first ANOVA, justified the inclusion of all children into one age group in the analyses in the main text.

However, we were surprised to find no age-group differences in the child sample and performed additional tests to assess age-related differences within the children, collapsing across age groups. We performed a linear regression to assess age-related heterogeneity using age as a continuous predictor. The dependent variable was fractional anisotropy (FA). Predictors included AGE as a continuous variable, PATH as a categorical variable, and the interaction between AGE and PATH. Sex was included as a dichotomous covariate of no interest. FA was significantly predicted by the model, $F(4, 106) = 2.910$, $p = 0.025$, with $R^2 = 0.102$ and $R_{adjusted} = 0.067$. The full model and resulting parameters were: Predicted FA = 1.941 - 0.27(AGE) - 2.94(PATH) + 0.004(AGE*PATH) + 0.158 (SEX). Each predictor was tested for significance using a one-sample *t*-test and a threshold of $p < 0.05$, using a Bonferroni correction for the 3 comparisons (i.e., the 3 predictors), $p_{Bonferroni} = 0.05/3 = 0.017$. There were no significant predictors after Bonferroni correction for multiple comparisons, all *p*s > 0.017. To further investigate, we conducted the linear regression with the interaction term removed: Predicted FA

= 1.012 - 0.016(AGE) - 0.003(PATH) + 0.163 (SEX). The overall model was significant, $F(3, 106) = 3.521$, $p = 0.018$. AGE was significant, $p = 0.004$, but PATH was not, $p = 0.961$, and SEX was not, $p = 0.319$. This supplemental analysis suggested, therefore, that FA in the tracts of interest to the current study changes significantly across the ages 5-8 years of age in our sample, but that the relationship between FA and age does not vary among pathways in this age range.

**Supplemental Analyses for "PVP tracts cluster with VHP tracts in children but not in adults"**

**Supplemental Analysis 2.1 Hemispheric cohesion**

MDS models for the left and right hemispheres independently were also tested. We compared the *SSD* values of the MDS fits to determine if hemisphere-specific models would be better suited for the data than models that included both hemispheres. We performed this comparison using the first two dimensions of the MDS results. The hemisphere specific models had either higher *SSD* values than both hemispheres in children (left: 0.43, right: 0.53, both: 0.35) or *SSD* values similar to both hemispheres in adults (left: 0.30, right: 0.27, both: 0.32), suggesting that MDS performed on both hemispheres produced a better or similar model fit than MDS performed on each hemisphere separately. For these reasons, the main analysis (see main text) was performed using both hemispheres together.

**Supplemental Analysis 2.2 Pathway cohesion**

To assess how well the tracts clustered into our three defined pathways (VHP, PVP, DHP; see **Supplemental Analysis 1.2** for a similar test), *k*-means clustering ($k = 3$) was applied to the first two dimensions of the MDS model. *k*-means was performed 10,000 times and the clustering result with the lowest Sum of Squared Distances (*SSD*) of the data points from the centroids was selected as the winning model (Arthur & Vassilvitskii, 2006; Seber, 2009). A silhouette score was calculated to estimate how closely the data points clustered around the centroids within each cluster in the winning model (Kaufman & Rousseeuw, 2009).

Results in children demonstrated that the VHP, PVP, and DHP tracts clustered into separated clusters (**Supplemental Figure 4a**). The model *SSD* was 0.366. The silhouette scores of the VHP cluster, the PVP cluster, and the DHP cluster were 0.709, 0.625, and 0.816, respectively, indicating that the clustering solution was appropriate. The similarity within tracts from the same pathway was greater than the similarity between tracts from different pathways in the child sample (**Supplemental Figure 4a**), suggesting that the VHP, PVP, and DHP can be treated as distinct pathways in our child sample (consistent with **Supplemental Analysis 1.2**).

Results in adults demonstrated that the MDLFspl tract clustered with the VHP tracts, that the pArc and TP-SPL clustered with the DHP tracts, and that the MDLFang was in a cluster of its own (**Supplemental Figure 4b**). The model *SSD* in adults was 0.293. The silhouette scores of clusters 1, 2, and 3 were 0.827, 0.469, and 1, respectively.

We show that white matter tracts cluster together into cohesive multi-tract pathways and development in the age range of our child sample (i.e., 5-8 years) happens at the pathway-level. The results of the MDS and *k*-means clustering analysis validated our pathways approach for the VHP, PVP, and DHP that was used in our primary analyses. We expected the ILF, IFOF, pArc, TP-SPL, MDLFspl, MDLFang, SLF1and2, and SLF3 to cluster into the VHP, PVP, and DHP pathways and we, therefore, set $k = 3$. Results support the pathway approach in our child sample but not necessarily in adults (**Supplemental Figure 4**). Finding that tracts cluster into the predicted pathways in the child sample but not in the adult sample suggests that the relationship among tracts within a pathway is more homogeneous in childhood than adulthood (see **Supplemental Analyses 1.2** and **2.2** for similar findings).

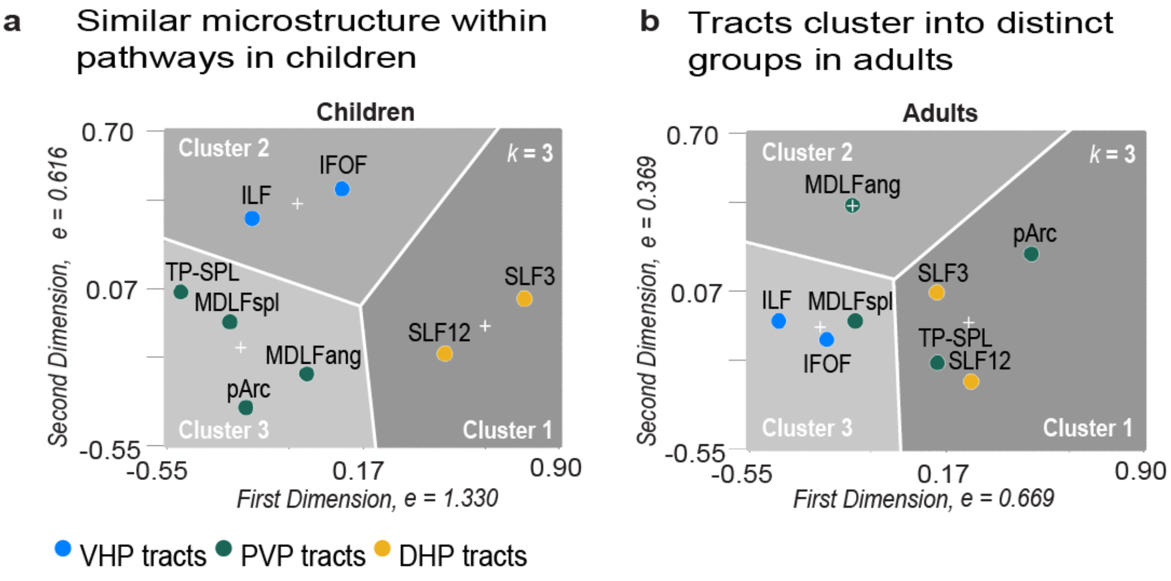


**Supplemental Figure 4. Separate clusters for VHP, PVP, and DHP tracts in children but not adults.** *k*-means clustering with $k = 3$ was performed to test tract membership in the ventral horizontal (VHP), posterior vertical (PVP), and dorsal horizontal (DHP) pathways. The analysis affirmed that the VHP, PVP, and DHP tracts clustered into their own clusters in the child data, but this was not the case for the adult data.

#### Supplemental Analysis 2.3 Bootstrap test of difference between PVP-VHP and PVP-DHP correlations

We performed an additional analysis to assess whether the correlation among PVP and VHP tracts was greater than the correlation among PVP and DHP tracts using a bootstrap test of the difference between the mean PVP-VHP and PVP-DHP correlations (see **Figure 4a**). Results showed that the PVP-VHP correlations were greater than the PVP-DHP correlations in children, $z = 2.145$, $p = 0.016$, but not in adults, $z = 0.073$, $p = 0.471$, consistent with the MDS and *k*-means results (**Figure 4c**).

#### Supplemental Analysis 2.4 Bootstrap test of difference between children and adults

We performed an additional analysis to assess whether the difference between the PVP-VHP and PVP-DHP correlations in children was statistically greater than the difference in adults, as would be suggested from the clustering results displayed in **Figure 4c**. We calculated the difference between PVP-VHP and PVP-DHP correlations in children and in adults. We then performed a bootstrap test of this difference between children and adults. Results showed that the difference between the PVP-VHP and PVP-DHP correlations in children was statistically greater than the difference in adults, $z$ = 1.841, $p$ = 0.033. This result supports the interpretation of the clustering solution provided by the visualization in **Figure 4c**: the PVP microstructure was similar to VHP microstructure in children, but not in adults. We note that this supplemental analysis uses the correlations among tract microstructural measures as the dependent variable and, therefore, cannot assess the spatial differences in the clustering solution.

**Supplemental analyses for "Performance on a visual perceptual task predicts PVP microstructure in children."**

Two analyses were performed to better understand if the results reported in the main manuscript could be explained by one hemisphere alone (**Supplemental Analysis 3.1**) and to see if a relationship with behavior could be found in pathways other than the PVP (**Supplemental Analysis 3.2**).

**Supplemental Analysis 3.1 Hemispheric cohesion**

We further tested for evidence of hemispheric asymmetry in the relationship between microstructure in the pArc and behavior. This analysis was important to ensure that combining the microstructure measures from the left and right hemispheres for each tract in the main analysis was appropriate. The left pArc was significantly predicted by behavior, but we did not find a significant relationship between the right pArc and behavior. For each hemisphere, we tested a linear model that predicted FA from the composite measures of LIT, VM, and FM that were used in the previous analyses. SEX was entered as a covariate of no interest.

*Left Hemisphere.* The model fit for the left pArc was significant, $F(5, 22) = 4.898$, $p = 0.006$, with $R^2 = 0.590$ and $R_{adjusted} = 0.470$. There were no significant predictors after Bonferroni correction, all $p$s > 0.017; however, the VM and FM composite scores trended towards significance with VM positively predicting FA in the pArc, $t(22) = 3.135$, $p = 0.006$, and FM negatively predicting FA in the pArc, $t(22) = 3.571$, $p = 0.002$.

*Right Hemisphere.* The model fit for the right pArc was significant, $F(5, 22) = 1.867$, $p = 0.153$, with $R^2 = 0.595$ and $R_{adjusted} = 0.165$, but there were no significant predictors, all $p$s > 0.017.

**Supplemental Analysis 3.2 Behavioral correlates of non-PVP pathways**

We performed the same multiple linear regression analysis described for the PVP for the other four pathways of interest: VOF, VHP, DHP, and FAT. None of the model fits were significant and there were no significant predictors.

*Vertical Occipital Fasciculus (VOF).* The model fit was not significant, $F(5, 22) = 0.624$, $p = 0.684$, with $R^2 = 0.155$ and $R_{adjusted} = -0.094$, and there were no significant predictors, all $p$s > 0.017.

*Ventral Horizontal Tracts (VHP).* The model fit was not significant, $F(5, 22) = 1.070$, $p = 0.411$, with $R^2 = 0.239$ and $R_{adjusted} = 0.016$, and there were no significant predictors, all $p$s > 0.017.

*Dorsal Horizontal Tracts (DHP).* The model fit was not significant, $F(5, 22) = 1.050$, $p = 0.421$, with $R^2 = 0.236$ and $R_{adjusted} = 0.011$, and there were no significant predictors, all $p$s > 0.017.

*Frontal Aslant Tract (FAT).* The model fit was not significant, $F(5, 22) = 1.670$, $p = 0.196$, with $R^2 = 0.329$ and $R_{adjusted} = 0.132$, and there were no significant predictors, all $p$s > 0.017.
